# Genetic insights into the social organisation of the Avar period elite in the 7^th^ century AD Carpathian Basin

**DOI:** 10.1101/415760

**Authors:** Veronika Csáky, Dániel Gerber, István Koncz, Gergely Csiky, Balázs G. Mende, Bea Szeifert, Balázs Egyed, Horolma Pamjav, Antónia Marcsik, Erika Molnár, György Pálfi, András Gulyás, Bernadett Kovacsóczy, Gabriella M. Lezsák, Gábor Lőrinczy, Anna Szécsényi-Nagy, Tivadar Vida

## Abstract

After 568 AD the Avars settled in the Carpathian Basin and founded the Avar Qaganate that was an important power in Central Europe until the 9^th^ century. Part of the Avar society was probably of Asian origin, however the localisation of their homeland is hampered by the scarcity of historical and archaeological data.

Here, we study mitogenome and Y chromosomal STR variability of twenty-six individuals, a number of them representing a well-characterised elite group buried at the centre of the Carpathian Basin more than a century after the Avar conquest.

The studied group has maternal and paternal genetic affinities to several ancient and modern East-Central Asian populations. The majority of the mitochondrial DNA variability represents Asian haplogroups (C, D, F, M, R, Y and Z). The Y-STR variability of the analysed elite males belongs only to five lineages, three N-Tat with mostly Asian parallels and two Q haplotypes. The homogeneity of the Y chromosomes reveals paternal kinship as a cohesive force in the organisation of the Avar elite strata on both social and territorial level. Our results indicate that the Avar elite arrived in the Carpathian Basin as a group of families, and remained mostly endogamous for several generations after the conquest.

## Introduction

The Carpathian Basin in East-Central Europe is generally regarded as the westernmost point of the Eurasian steppe, and as such, its history was often influenced by the movements of nomadic people of eastern origin. After 568 AD, the Avars settled in the Carpathian Basin and founded their empire which was a powerful player in the geopolitical arena of Central and Eastern Europe for a quarter of a millennium^1,2^.

The supposed Asian origin of the Avars appeared as early as the 18^th^ century. Since then various research approaches emerged indicating different regions as their home of origin: i.e. Central or East-Central Asia (see SI chapter 1b for explanation of this geographic term). This debate remained unsolved, however a rising number of evidences points towards the latter one^1,2^.

The history of the Avars is known from external, mainly Byzantine written accounts of diplomatic and historical character focusing on certain events and important people for the Byzantine Empire. As an example, the description of a Byzantine embassy in 569-570 visiting the Western Turkic Qaganate in Central Asia, claimed that their ruler complained about the escape of his subjects, the Avars^2–4^.

The linguistic data concerning the Avars are limited to a handful of personal names and titles (Qagan, Bayan, Yugurrus, Tarkhan, etc.) mostly of East-Central Asian origin, known from the same Byzantine written accounts. The available evidence is not sufficient for defining the affiliation of the Avars’ language, however the scarce remains suggest Proto-Mongolian, Proto-Turkic and/or a still undefined Central Asian or Siberian language^1,2,5^.

New elements appeared with the Avars in the archaeological heritage of the Carpathian Basin that shared common characteristics with Eurasian nomadic cultures. These phenomena are even more emphasised in the burials of the Avar period elite group composed of only a dozen graves^6,7^.This group of lavishly furnished burials -the focus of our study-is located in the Danube-Tisza Interfluve (central part of the Carpathian Basin) and is dated to the middle of the 7^th^ century (Fig. 1). They are characterised by high-value prestige artefacts such as gold- or silver-plated ring-pommel swords, gold belt-sets with pseudo-buckles and certain elements of precious metal tableware (see SI chapter 1c, Fig.2). The concentration of these burials can, in all likelihood, be linked to leaders of the early Avar polity and the Qagan’s military retinue^6,7^. The Avar-period material culture shows, how this ruling elite remained part of the connection network that is the Eurasian steppe, even generations after settling in the Carpathian Basin (SI chapter 1c).

**Figure 1.**
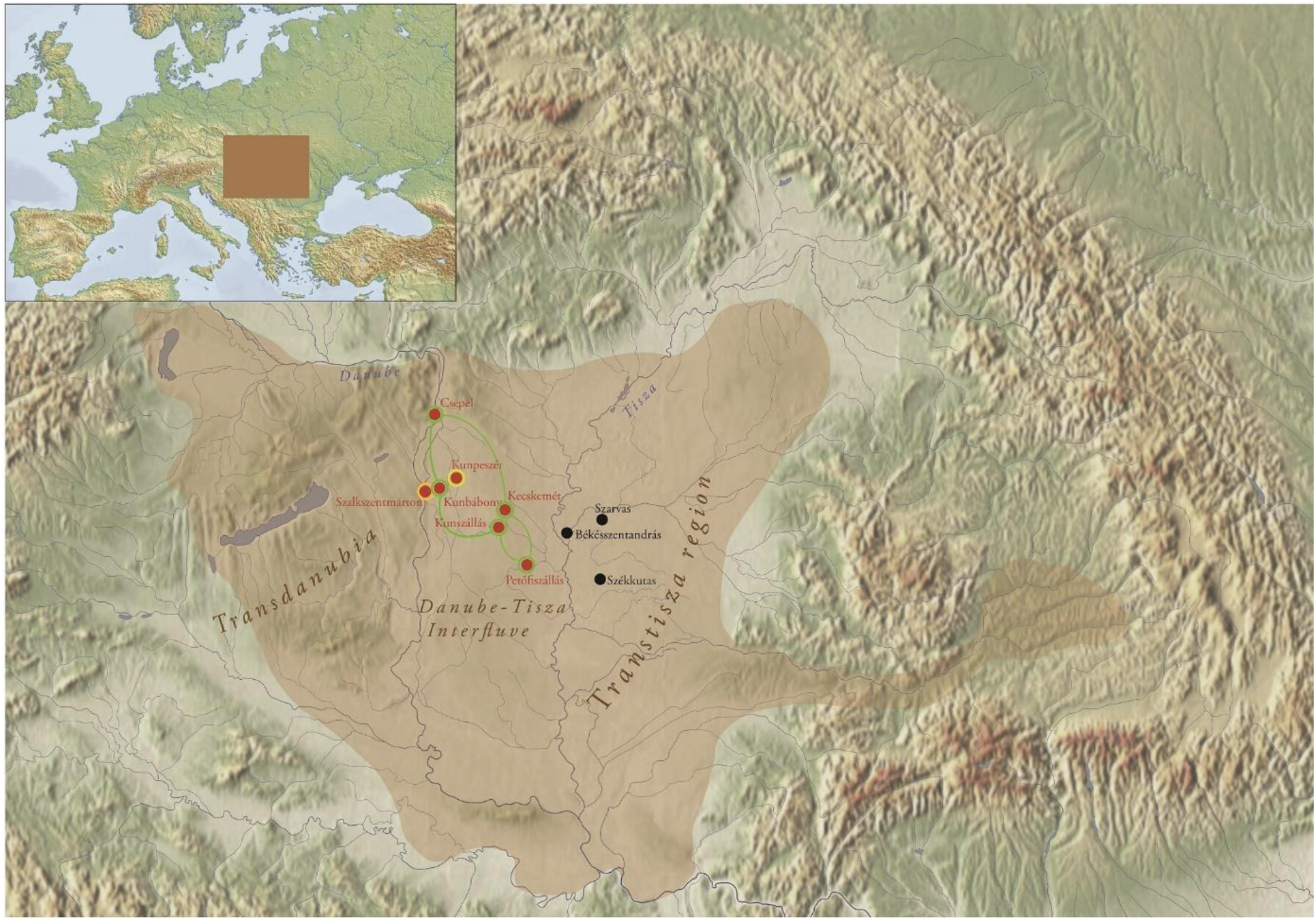
Territory of the early Avar Qaganate and the location of the investigated sites in the Carpathian Basin. The investigated sites of the Kunbábony group (7^th^ century) are marked with red, 7^th^-8^th^ century supplementary sites are marked with black dots. Yellow and orange circles indicate the detection of Y chromosomal N-Tat haplotype I and III respectively. Green circles and lines indicate the occurrence of shared N-Tat haplotype II in five burial sites of the Avar elite. Brown shade indicates the territory of the early Avar Qaganate. The map of the Carpathian Basin is owned by the IA RCH HAS, and was modified in Adobe Illustrator CS6. The map of Europe shown in the upper left corner, licensed under CC BY 4.0, was downloaded from MAPSWIRE (https://mapswire.com/europe/physical-maps/).

**Figure 2.**
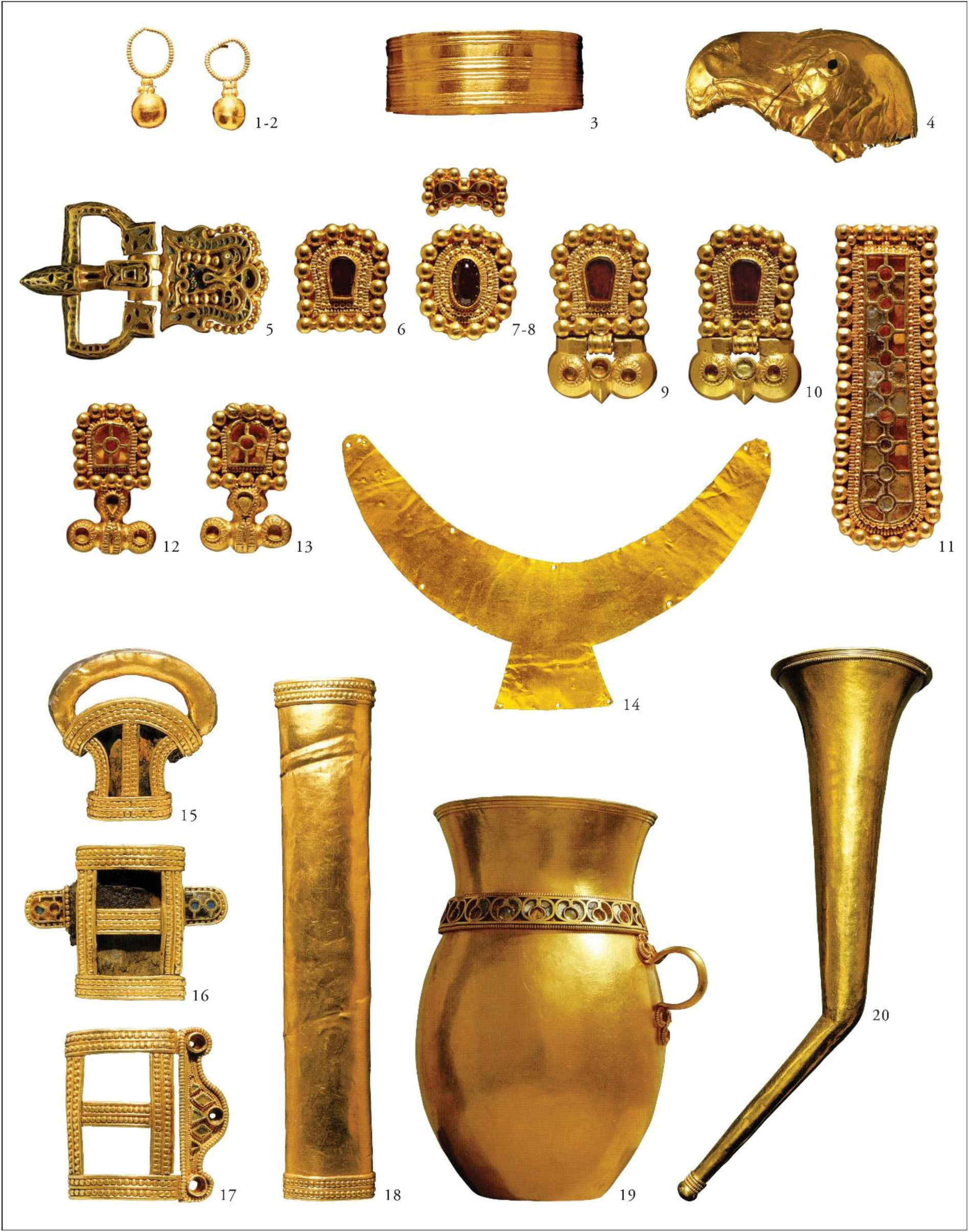
A selection of grave goods from the burial at Kunbábony. The burial of an adult man at Kunbábony (AC2) contained 2.34 kilograms of gold in form of weaponry covered with precious metal foils, ornamented belt sets with so-called pseudo buckles and drinking vessels. The funerary attire and the grave goods are understood as elements of the steppe nomadic material culture of the period. The technological details and the decoration however suggest a culturally heterogeneous origin. Presented objects: 1-2. earrings; 3. armring; 4. eagle head-shaped end of a sceptre or horsewhip; 5-13. elements of the belt with the so-called pseudo buckles (9-10.); 14. crescent-shaped gold sheet; 15-18. sword fittings; 19. jug; 20. drinking horn. Pictures were first published in H. Tóth & Horváth^15^.

The Carpathian Basin witnessed population influxes from the Eurasian Steppe several times, which are genetically poorly documented. The earliest such migration was that of the *Yamnaya* people in the 3^rd^ millennium BC^8^. Further eastern influxes reached the Carpathian Basin with Iron Age Scythians, the Roman Age Sarmatians and with the Huns in the 5^th^ century. The few analysed Scythian samples from Hungary had relatively increased European farmer ancestry and showed no signs of gene flow from East-Central Asian groups^9^. The Sarmatians and Huns from Hungary have not yet been studied.

Besides influxes from the east, the Carpathian Basin witnessed population movements from the north as well. The Lombards for e.g., who directly preceded the Avars in Transdanubia (today’s Western Hungary), showed Central and North European genomic ancestry in recent studies^10,11^. Few ancient DNA studies have focused on the Avars, and these studies analysed only the control region of the mitochondrial DNA (mtDNA). One research focused on a 7^th^-9^th^ century Avar group from the south-eastern part of the Great Hungarian Plain (Alföld) of the Carpathian Basin^12^. Their maternal gene pool showed predominantly Southern- and Eastern-European composition, with Asian elements presenting only 15.3% of the variation. Another recent study of a mixed population of the Avar Qaganate from the 8^th^-9^th^ centuries from present-day Slovakia showed a miscellaneous Eurasian mtDNA character too, with a lower frequency (6.52%) of East Eurasian elements^13^.

Here we study 26 Avar period individuals, who were excavated at ten different sites (found in small burial groups or single burials). Seven out of ten sites are located in the Danube-Tisza Interfluve^7,14,15^, while three are located east of the Tisza river where a secondary power centre can be identified in the 7^th^ century^16^ (Fig. 1). The primary focus of the sample selection was to target all available members (eight individuals) of the highest elite Avar group from the Danube-Tisza Interfluve complemented by other individuals from the Tisza region (see Materials and SI chapter 1a).

Our main research questions concern the origin and composition of this ruling group of the Avar polity. Was it homogeneous or heterogeneous? Is it possible to identify a migration and if yes what can we tell about its nature? Were maternal and paternal lineages of similar origin? Did kinship groups play a role in the organisation of this elite strata?

Using whole mitogenome sequencing, and Y chromosomal short tandem repeat (Y-STR) analyses, our current research focused on the uniparental genetic diversity of the leading group of the Avar period society from the 7^th^ century AD.

## Results

### Primary observations

We sequenced mitochondrial genomes, using a hybridisation capture method, of 25 Avar period individuals (42x average coverage, the 26. sample was tested only for Y-STR, see Table S1 for details). Osteological sex determination was checked by shallow shotgun sequencing data. The studied Avar group composed of 18 males and 8 females.

The mitochondrial genome sequences can be assigned to a wide range of the Eurasian haplogroups with dominance of the Asian lineages, which represent 69.5% of the variability: four samples belong to Asian macrohaplogroup C (two C4a1a4, one C4a1a4a and one C4b6); five samples to macrohaplogroup D (one by one D4i2, D4j, D4j12, D4j5a, D5b1), and three individuals to F (two F1b1b and one F1b1f). Each haplogroup M7c1b2b, R2, Y1a1 and Z1a1 is represented by one individual. One further haplogroup M7 (probably M7c1b2b), was detected (sample AC20); however, the poor quality of its sequence data (2.19x average coverage) did not allow further analysis of this sample.

European lineages (occurring mainly among females) are represented by the following haplogroups: H (one H5a2 and one H8a1), one J1b1a1, two T1a (two T1a1), one U5a1 and one U5b1b (Table S1). One further T1a1b sample (HC9) came from a distinct cultural group of the Avar society, and therefore was not included to the comparative analyses on the Avar elite.

The Y-STR analyses of 17 males give evidence on a surprisingly homogeneous Y chromosomal composition (Table S1). Y chromosomal STR profiles of 14 males could be assigned to haplogroup N-Tat (also N1a1-M46, see Methods and Table S1). N-Tat haplotype I was found in four males from Kunpeszér with identical alleles on at least nine loci. The full Y-STR haplotype I, reconstructed from AC17 with 17 detected STRs, is rare in our days. Only nine matches were found among 205,059 haplotypes in YHRD database, such as samples from the Ural Region, Northern Europe (Estonia, Finland), and Western Alaska (Yupiks). We performed Median Joining (MJ) network analysis using 162 N-Tat haplotypes with ten shared STR loci (Fig. 3, Table S9). All modern N-Tat samples included in the network had derived allele of L708 as well. Haplotype I (Cluster 1 in Fig. 3) is shared by eight populations on the MJ network among the 24 identical haplotypes. Cluster 1 represents the founding lineage, as it is described in Siberian populations^17^, because this haplotype is shared by the most populations and it is more diverse than Cluster 2.

**Figure 3.**
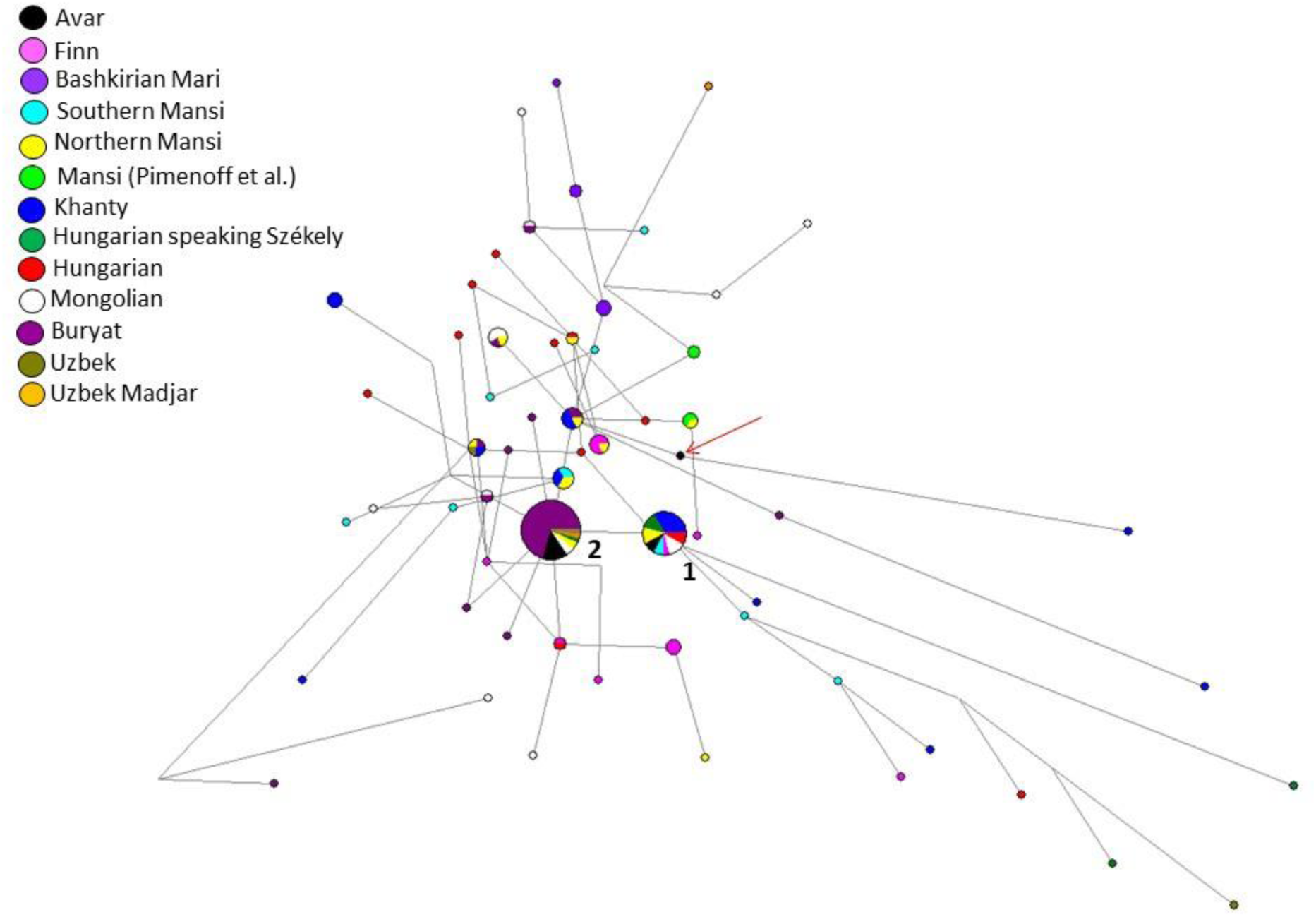
Median Joining network of 162 N-Tat Y-STR haplotypes. Allelic information of ten Y-STR loci were used for the network. Only those Avar samples were included, which had results for these ten Y-STR loci. The founder haplotype I (Cluster 1) is shared by eight populations including three Mongolian, three Székely, three northern Mansi, two southern Mansi, two Hungarian, eight Khanty, one Finn and two Avar (AC17, AC26) chromosomes. Haplotype II (Cluster 2) includes 45 haplotypes from six populations studied: 32 Buryats, two Mongolians, one Székely, one Uzbek, one Uzbek Madjar, two northern Mansi and six Avars (AC1, AC12, AC14, AC15, AC19 and KSZ 37). Haplotype III (indicated by a red arrow) is AC8. Information on the modern reference samples is seen in Table S9.

Nine males share N-Tat haplotype II (on a minimum of eight detected alleles), all of them buried in the Danube-Tisza Interfluve (Table S1). We found 30 direct matches of this N-Tat haplotype II in the YHRD database, using the complete 17 STR Y-filer profile of AC1, AC12, AC14, AC15, AC19 samples. Most hits came from Mongolia (seven Buryats and one Khalkh) and from Russia (six Yakuts), but identical haplotypes also occur in China (five in Xinjiang and four in Inner Mongolia provinces). On the MJ network, this haplotype II is represented by Cluster 2 and is composed of 45 samples (including 32 Buryats) from six populations (Fig. 3). A third N-Tat lineage (type III) was represented only once in the Avar dataset (AC8), and has no direct modern parallels from the YHRD database. This haplotype on the MJ network (see red arrow in Fig. 3) seems to be a descendent from other haplotype cluster that is shared by three populations (two Buryat from Mongolia, three Khanty and one Northern Mansi samples). This haplotype cluster also differs one molecular step (locus DYS393) from haplotype II.

We classified the Avar samples to downstream subgroup N-F4205 within the N-Tat haplogroup, based on the results of ours and Ilumäe et al.^18^ and constructed a second network (Fig. S4). The N-F4205 network results support the assumption that the N-Tat Avar samples belong to N-F4205 subgroup (see SI chapter 1d for more details).

Based on our calculation, the age of accumulated STR variance (TMRCA) within N-Tat lineage for all samples is 7.0 kya (95% CI: 4.9 - 9.2 kya), considering the core haplotype (Cluster 1) to be the founding lineage. (See detailed results on the N-Tat and N-F4205 haplotypes in the SI chapter 1f.) Y haplogroup N-Tat was not detected by large scale Eurasian ancient DNA studies^9,19^ but it occurs in late Bronze Age Inner Mongolia^20^ and late medieval Yakuts^21^, among them N-Tat has still the highest frequency^22^.

Two males (AC4 and AC7) from the Transtisza group belong to two different haplotypes of Y-haplogroup Q1. Both Q1a-F1096 and Q1b-M346 haplotypes have neither direct nor one step neighbour matches in the worldwide YHRD database. A network of the Q1b-M346 haplotype shows that this male had a probable Altaian or South Siberian paternal genetic origin (Fig. S5).

### Possible kinship connections in the cemetery at Kunszállás

We detected two identical F1b1f mtDNA haplotypes (AC11 female and AC12 male) and two identical C4a1a4 haplotypes (AC13 and AC15 males) from the same cemetery of Kunszállás; these matches indicate possible maternal kinship of these individuals. Further pair is AC9 female and AC14 male, who shared the same T1a1 mtDNA lineage.

The detected Y chromosomal lineages probably all belong to one shared N-Tat haplotype in Kunszállás (AC12, AC13, AC14, AC15), which indicate that it was a cemetery used by both maternally and paternally closely related individuals.

### Comparative analyses of the ancient dataset

The elite group originating mainly from the Danube-Tisza Interfluve does not exhibit a genetic connection to the previously investigated small Avar period population from southeast Hungary^12^, because the latter shows predominantly Eastern European maternal genetic composition. This result is comparable with the archaeological records, i.e. this Avar population buried the deceased in catacomb graves, following Eastern European traditions.

One sample in our dataset (HC9) comes from this population, and both his mtDNA (T1a1b) and Y chromosome (R1a) support Eastern European connections. The observed within-Avar genetic differences correlate well with the cultural and anthropological differences of this group and demonstrate the heterogeneity of the Avar population.

We also find that the Avar elite group is genetically different from the 6^th^ century Lombard period community of Szólád in Transdanubia^10^, which has more genetic affinity to other ancient European populations (Fig. 4).

**Figure 4.**
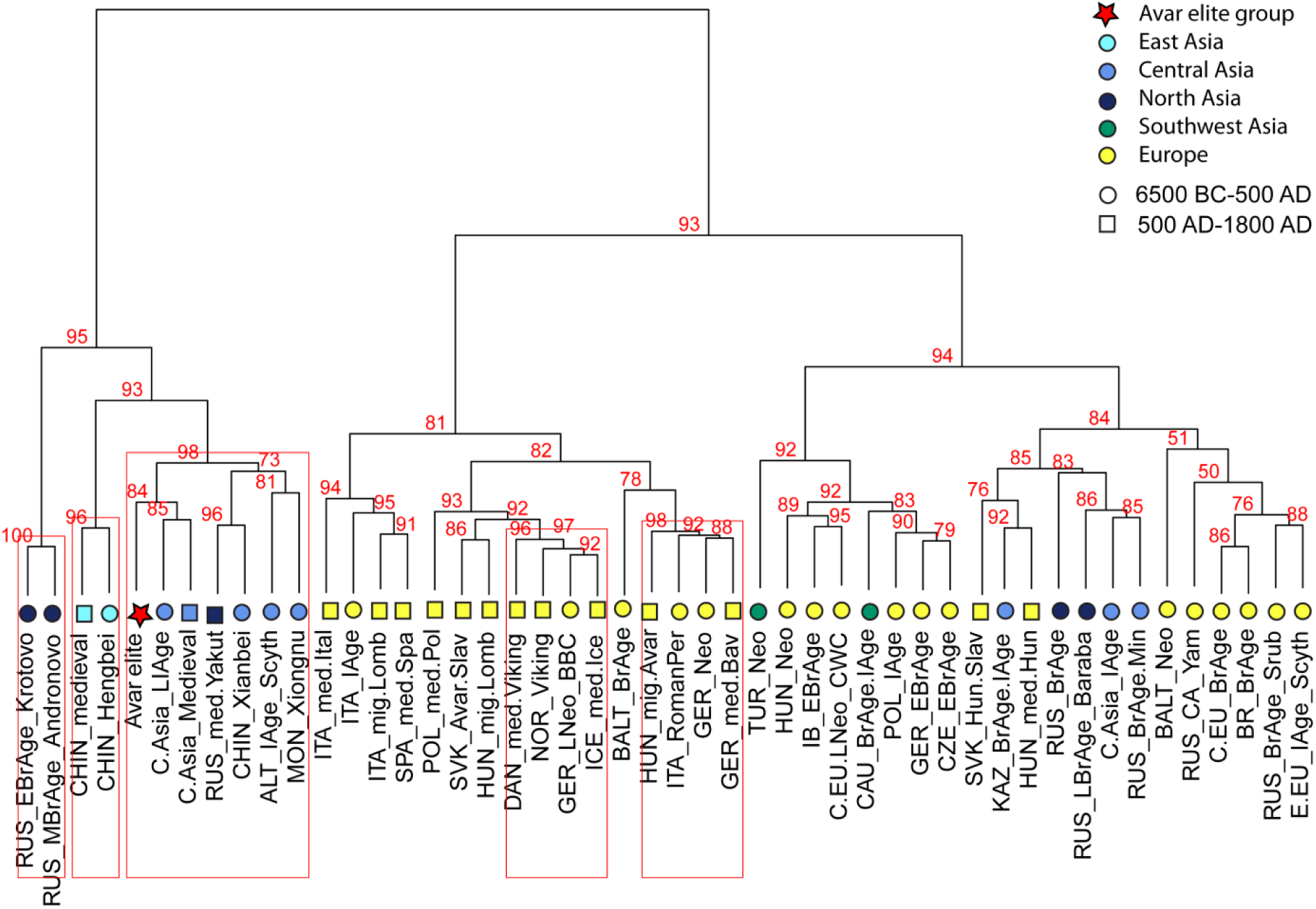
Ward type clustering of 48 ancient populations. The Ward type clustering shows separation of Asian and European populations. The Avar elite group (AVAR) is situated on an Asian branch and clustered together with Central Asian populations from Late Iron Age (C-ASIA_LIAge) and Medieval period (C-ASIA_Medieval), furthermore with Xiongnu period population from Mongolia (MON_Xiongnu), Xianbei period of China (CHIN_Xianbei), medieval Yakuts (RUS_med.Yakut) and Altaian Scythians (ALT_IAge_Scyth). P values are given in percent as red numbers on the dendogram, where red rectangles indicate clusters with significant p values. The abbreviations and references are presented in Table S2.

Comparing this early Avar period elite with later datasets from the Carpathian Basin, only a few connections are observable. The mixed population of the Avar Qaganate dated to the 8^th^- 9^th^ centuries^13^ does not show affinities to the studied group.

The overall mtDNA composition of the Avar elite group and the 9^th^-12^th^ century populations of the Carpathian Basin differ significantly, population continuity is not observable. The T1a1b mtDNA phylogenetic tree contains one individual from the Hungarian conquest period (sample Karos III/14^23^) with identical sequence to the Avar HC9, which might indicate the genetic continuity of certain maternal lineages between the 7^th^ and 9^th^-10^th^ centuries. Some further haplogroup-level matches exist between the ancient Hungarians^24^ and Avars, but these do not mean close phylogenetic relationships. On the other hand, Y chromosomal N-Tat haplotypes show that certain paternal lineages could have continuity among the Avars and Early Hungarians (see SI chapter 1f).

A possible continuity of the Avar population should be studied on larger dataset covering the entire spectrum of the Avar society.

In the comparative analyses we included ancient mtDNA data from whole Eurasia, especially focusing on geographically or chronologically relevant sample sets from the Carpathian Basin, Central and East Asia.

We performed Principal Component Analysis (PCA) with the Avar dataset using haplogroup frequencies of another 47 ancient groups (Table S2, Figs. S6a-b). The Avar elite shows affinities to some Asian populations: they are close to 15^th^-19^th^ centuries Yakuts from East-Siberia and to two ancient populations from China along PC1 and PC2, while along PC3, the Avars are near to South Siberian Bronze Age populations, which is possibly caused by high loadings of the haplogroup vectors T1 and R on PC3. The strict separation of Asian and European populations is also displayed on the Ward-type clustering tree. Here the Avar elite is located on an Asian branch of the tree and clustered together with Iron Age and medieval Central and East-Central Asian sample sets (Fig. 4).

Because whole mitochondrial genome datasets of ancient populations are still scarce (especially east of the Altai), we applied a smaller reference dataset (n=932) in the genetic distance calculations using full mitogenomes (see Fig. S9).

The Avar group shows significant genetic distance (p <0.05) from most ancient populations. Only two groups from Central Asia have non-significant differences from the Avar elite: one group containing Late Iron Age samples (originating from the Late Iron Age and Hun period from the Kazakh Steppe and the Tian Shan) (F_ST_ = −0.00116, p = 0.42382), and a group of Medieval samples from the Central Asian Steppe and the Tian Shan^9^ (F_ST_ = 0.00650, p = 0.26839, Table S4). These groups however contain scattered samples from large geographic area and period, therefore only limited inference can be drawn. Building of these large Central Asian sample pools was necessitated by the small number of samples per cultural groups in the reference studies from Asia.

The multidimensional scaling (MDS) plots based on linearised Slatkin F_ST_ values (Tables S4 and S5) of 26 ancient groups does not show a clear chronological or geographical grouping; however, Asia and Europe are separated. The Avar elite group is close to Central-Asian groups from the Late Iron Age and Medieval period^9^ in accordance with the individual F_ST_ results (Fig. 5).

**Figure 5.**
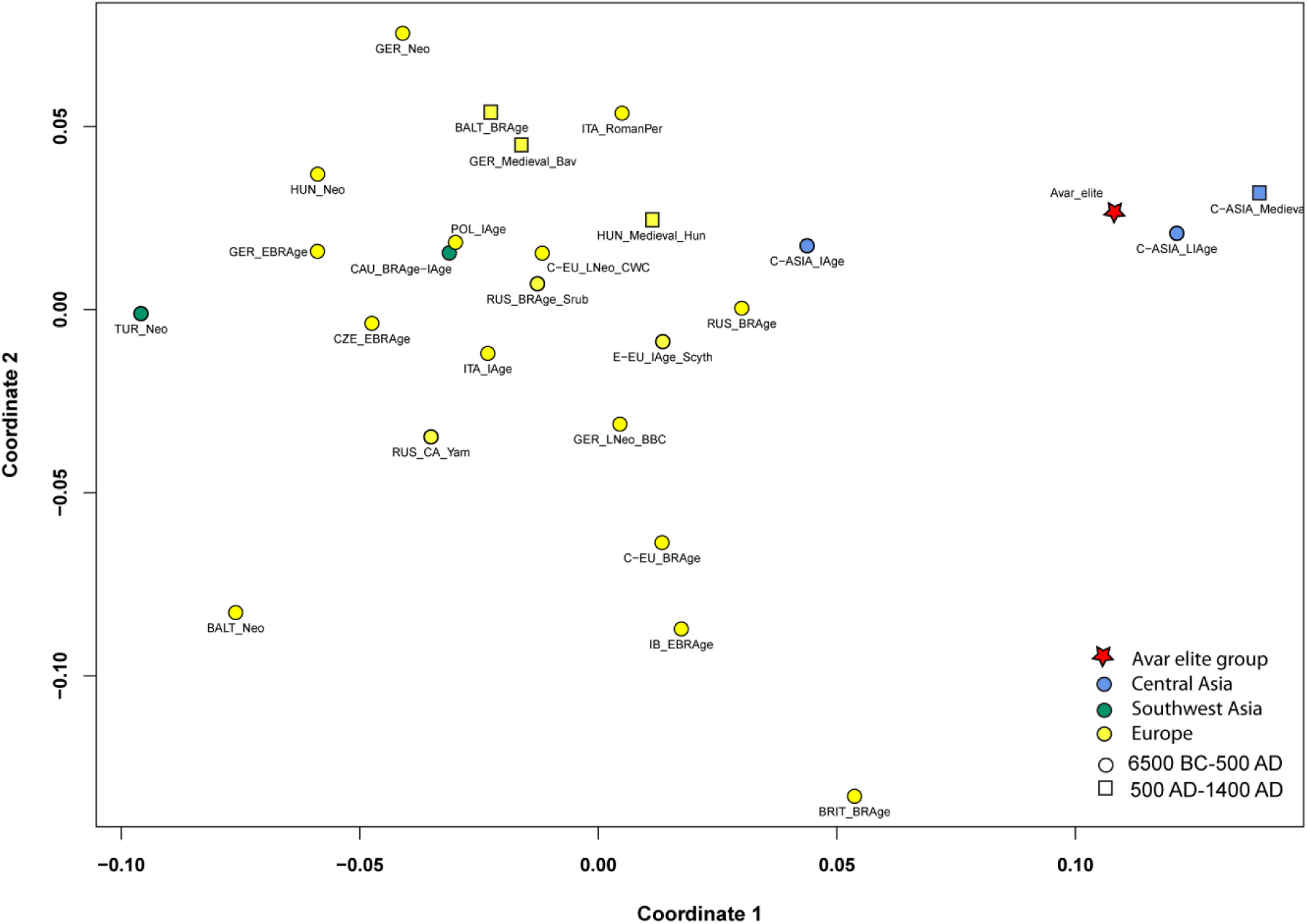
MDS with 26 ancient populations. The multidimensional scaling plot is based on linearised Slatkin F_ST_ values that were calculated based on whole mitochondrial sequences (stress value is 0.1669). The MDS plot shows the connection of the Avar elite group to the Central-Asian populations of the Late Iron Age (C-ASIA_LIAge) and Medieval period (C-ASIA_Medieval) along coordinate 1 and coordinate 2, which is caused by small genetic distances between these populations. The European ancient populations are situated on the left part of the plot. The F_ST_ values, abbreviations and references are presented in Table S4.

### Summary of the modern East-Eurasian maternal genetic affinities of the Avar elite group

Although DNA composition of modern populations can only give us indirect information about past populations, the lack of ancient Asian reference data leads us to use modern populations as proxy to ancient peoples in the phylogeographic analyses.

We performed PCA and MDS with modern mitogenome datasets (Table S3, Figs. 6, S7) and separately counted and constructed Neighbour Joining (NJ) phylogenetic trees of the 16 mtDNA haplogroups detected (see Table S6, Methods, SI, Figs. S10a-o). The NJ trees of certain haplogroups provide evidence of the phylogenetic connection of the 16 Avars samples with individuals from Asian populations.

**Figure 6.**
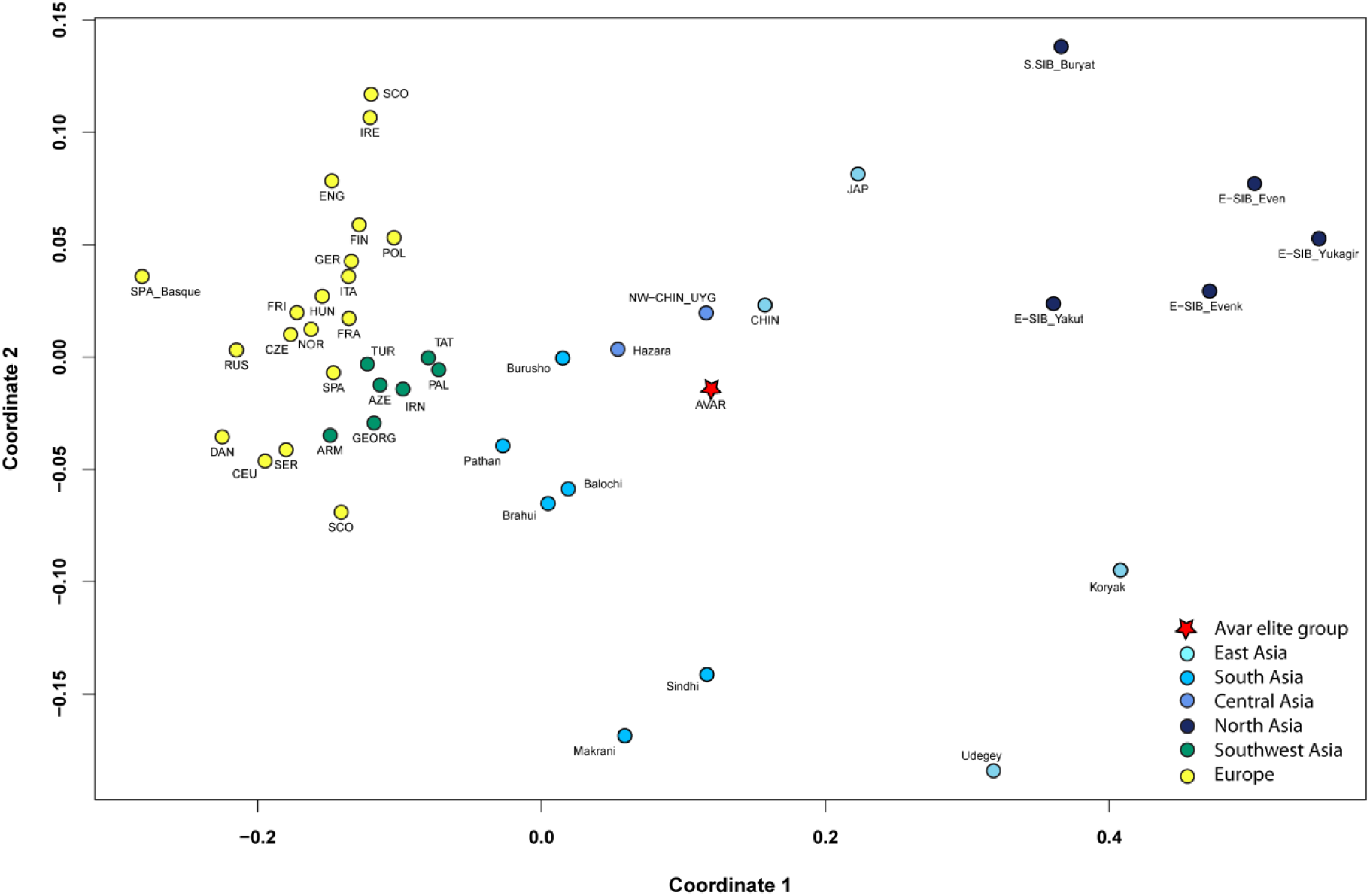
MDS with the 44 modern populations and the Avar elite group. The multidimensional scaling plot is displayed based on linearised Slatkin F_ST_ values calculated based on whole mitochondrial sequences (stress value is 0.0677). The MDS plot shows differentiation of European, Near-Eastern, Central- and East-Asian populations along coordinates 1 and 2. The Avar elite (AVAR) is located on the Asian part of plot and clustered with Uyghurs from Northwest-China (NW-CHIN_UYG) and Han Chinese (CHIN), as well as with Burusho and Hazara populations from the Central-Asian Highland (Pakistan). The F_ST_ values, abbreviations and references are presented in Table S5.

Modern East-Siberian populations, namely Yakuts and Nganasans are close to the Avar elite based on their haplogroup composition (Figs. S7-S8, Table S3). Phylogenetic connections to the Yakuts and Nganasans as well as to further East-Siberian individuals (Evenks and Tungusic people) are presented in C4a1a, D4i, D4j, F1b1, Y1a and Z1a NJ trees (Figs. 7 and S10a, S10c, S10e, S10h, S10o). The Yakuts had an East-Central Asian origin^25^. Shared lineages with the Avars might refer to the relative proximity of their homelands, or admixture of the Yakuts with Mongolians before their migration to the north. The mtDNA results of Yakuts show a very close affinity with Central Asian and South Siberian groups, which also suggests their southern origin^22^.

**Figure 7.**
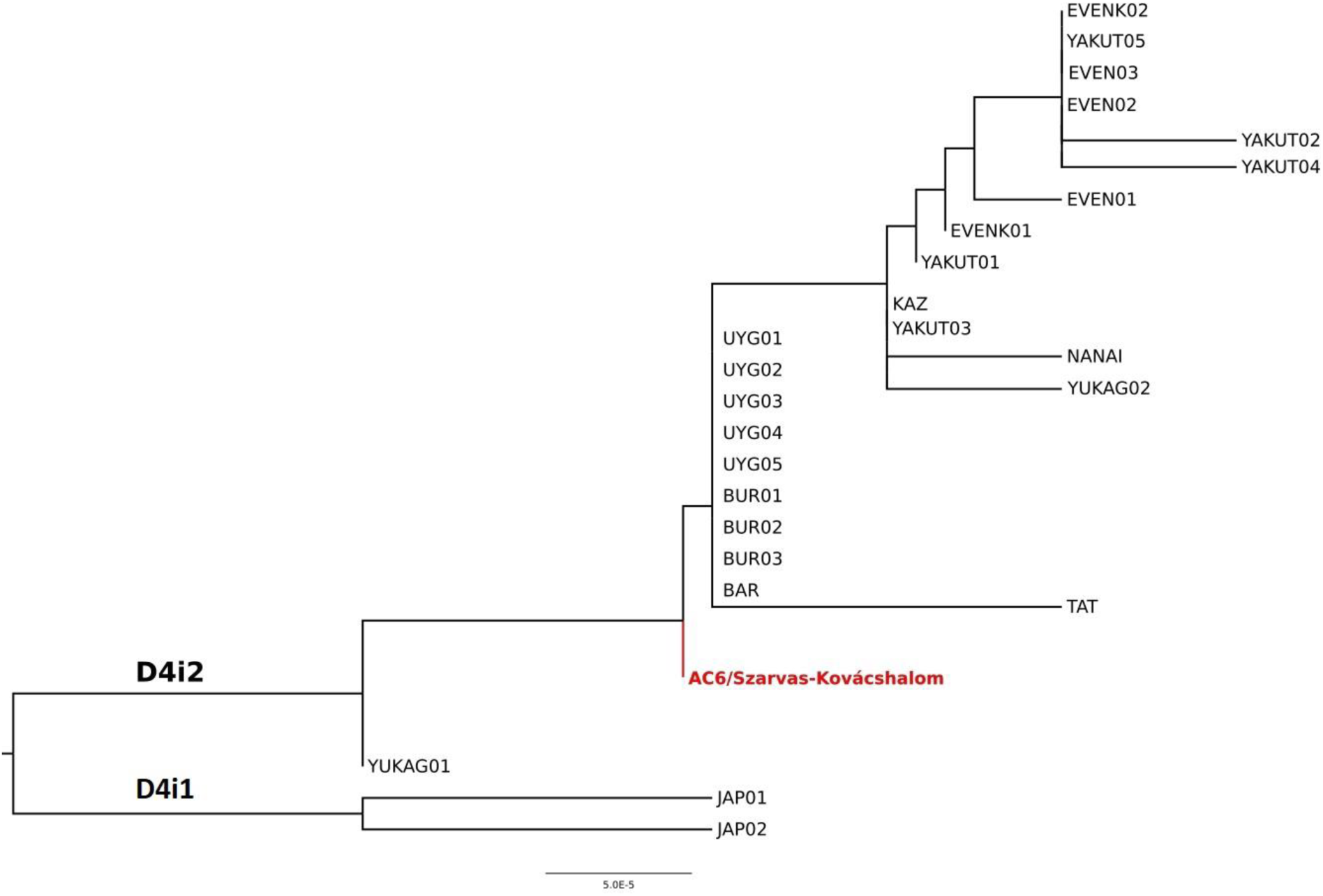
Phylogenetic tree of D4i2 sub-haplogroup. Phylogenetic tree of D4i2 sub-haplogroup shows AC6 to be the mitochondrial founder of most of the other D4i2 lineages from in East-Central and North Asia, which indicates a close shared maternal ancestry between the populations represented by these individuals. The references of individuals displayed on the tree are presented in Table S6.

Genetic connection with Russian Trans-Baikalian Mongolian-speaking Buryats and Barguts of the Avars is displayed on C4a1a, D4i and D4j phylogenetic trees (Figs. 7, S10a, S10c). The Buryats also stay on one branch with the Avars on the Ward-type clustering tree (Fig. S6). Furthermore, the Buryats appear on C4b, F1b1 and Y1a phylogenetic trees as well (Figs. S10b, S10e, S10h). Derenko et al. recently summarised the genetic research of the Buryats, who show connections to Chinese and Japanese but also to Turkic and Mongolic speaking populations^26^. Yunusbayev et al. concluded based on genome wide genotype data that Tuvinians, Buryats and Mongols are autochthonous to their current southern Siberian and Mongolian residence^27^. The Buryats represent a population that did not migrate much in the last millennia; therefore, they can be a good proxy for the Medieval population of South Siberia. Unfortunately, modern whole mitogenomic data are underrepresented from the East-Central Asian region (e.g. Mongolia), which region was (according to historical records) an important source of early Medieval nomadic migrations.

The genetic connection of Avar period elite group with modern Uyghurs from Northwestern-China (Xinjiang, Turpan prefecture)^28^ is shown by the detected low genetic distance between the Avar elite group and modern Uyghur individuals compared to the other 43 modern populations (Table S5, Fig. 6). The Uyghurs are relatively near to the Avar period elite on the PCA plots and on the Ward-clustering tree (Figs. S7-S8, Table S3). The NJ trees of haplogroups C4b, D4i, D4j, D5b, F1b1, M7c1b2, R2, Y1a and Z1a also give evidence of the phylogenetic connections to the modern Uyghurs (Figs. 7 and S10b-h, S10o). However, it is important to emphasise that this population is not the descendent of the Medieval *Uighur* Empire, since modern Uyghurs gained their name only during the 20^th^ century^29^.

The genetic distance is small between the investigated Avar elite and some modern-day ethnic groups from the Central-Asian Highlands (lying mostly in the territory of Afghanistan and Pakistan) (Table S5)^30^, the connections of which are shown on the MDS plot (Fig. 6) and on the haplogroup R2 tree as well (Fig. S10g). Interestingly, the Hazara population, living mostly in Afghanistan and Pakistan today, probably has a Mongolian origin^31^. Further Central-Asian individuals from the Pamir Mountains show phylogenetic connection with Avars on the D4j, R2 trees, and interestingly also on the European T1a1b tree (Figs. S10c, S10g, S10n).

The Central-Asian Kazakhs and Kyrgyz cluster together with the Avar group on PCA plots and clustering tree (Figs. S7-8, Table S3). Unfortunately, they cannot be presented on the MDS plot because of the absence of available population-level whole mitogenomic data. However, one modern Kazakh individual with the D4i haplogroup shares a common ancestor with an Avar period individual AC6 (Fig. 7).

Caucasian genetic connection is presented only by one sample on the phylogenetic tree of haplogroup H8a, where the AC17 sample from the Avar period is situated on one branch that also contains ancient and modern Armenians (Fig. S10j).

## Discussion

In 568 AD the Avars arrived in the Carpathian Basin, which was inhabited in the 6^th^ century by mixed Barbarian and Late Antique (Romanised) groups^32^. The Avar Qaganate can be regarded as composition of heterogeneous groups regardless of the linguistic, cultural or ethnic affiliations^2^. The highest social stratum however shows a homogeneous cultural and anthropological character. The historical sources suggest that this group introduced titles and institutions of a nomadic state in the Carpathian Basin^1,2^.

### Genetic data on the origin of the Avar elite

The paternal genetic data of the studied Avar group is very homogeneous compared to the maternal gene pool, and mostly composed of N-Tat haplotypes. Two males buried in the cemetery of Kunpeszér, have an N-Tat Y-STR haplotype I that has direct parallels to Buryat, Mongolian, Uzbeks, Hungarian speakers and Mansi (Figs. 3, S4). The second N-Tat haplotype (haplotype II) could signalize shared common genetic history of Avars with ancestors of Mongolians and Uralic populations (Figs. 3, S4). According to Ilumäe et al. study^18^, the frequency peak of N-F4205 (N3a5-F4205) chromosomes is close to the Transbaikal region of Southern Siberia and Mongolia, and we conclude that most Avar N-Tat chromosomes probably originated from a common source population of people living in this area, completely in line with the results of Ilumäe et al.^18^.

In the Transtisza region no N-Tat haplogroup appears, but two different Q1a and Q1b Y-STR profiles are detected at Szarvas-Kovácshalom. They do not have direct haplotype parallels, and these Q sub-haplogroups have a wide distribution in Eurasia. A network of the Q1b-M346 haplotype however shows that AC7 had a probable Altaian or South Siberian paternal genetic origin (Fig. S5).

The maternal gene pool of the investigated Avar elite group is more complex, it contains both Western and Eastern Eurasian elements; nevertheless, Eastern Eurasian maternal lineages dominate the diverse spectrum in 69.5%. Only loose connections are detectable between the Avar elite and the available mtDNA data of ancient populations in Eurasia, with the highest affinities to Central and East-Central Asian ancient populations. The comparisons are encumbered by the geographically and chronologically scattered nature of the available ancient whole mitogenomes (Fig. S9). The comparative mitogenomic dataset is especially insufficient from the East-Central Asian territories. There is a sole sequenced genome from Mongolia (Khermen Tal) dated to the 5^th^ century that belongs to mtDNA haplogroup D4b1a2a1, whose frequency had probable increase in the Asian population ca. 750 years ago^33^. All the D4b1a and the D4i2, D4j, D5b groups (the latter three detected among our samples) are common in the modern populations of East-Central Asia (Table S6). This region was ruled by the Rouran Qaganate between the 4^th^ and the 6^th^ century AD. Based on historical research this area could have been one of the source regions of the Avar migration^2,34^(see SI chapter 1c). Further DNA data from Central and East Asia are needed to specify the ancient genetic connections; however, genomic analyses are also set back by the state of archaeological research, i.e. the lack of human remains from the 4^th^-5^th^ century Mongolia, which would be a particularly important region in the study of the Avar period elite’s origin^35^.

Due to the lack of ancient reference data we also compared the maternal and paternal genetic data of the investigated elite group to modern Eurasian populations. The results support the East-Central Asian genetic dominance in the genetic composition of the Avar elite group. The Avar period group shows low genetic distances and close phylogenetic connections to several modern East Eurasian populations. Phylogeographic data on individual mtDNA lineages points toward East-Central Asian populations such as Uyghurs and Buryats. Further genetic connections of the Avars to modern populations living to southwest (Hazara) and north (Yakuts, Tungus, Evenk) of East-Central Asia probably indicate common source populations.

The archaeological heritage of the Avar elite does not contradict our results. Certain artefacts found in the burials of the Kunbábony group point to eastern cultural connections, but a more precise definition is hindered by their different distribution patterns (see SI chapter 1c). Ring-pommel swords covered with golden or silver sheets were used as prestige goods from the Carpathian Basin as far as the Altai Region, or even China, Korea and Japan (Fig. S1)^7^. The crescent-shaped gold sheet from Kunbábony interpreted as an insignia could indicate a more symbolic connection towards Mongolia and Northern China (Fig. S3)^35–37^. The presence of these artefacts is not necessarily connected to the migration of individuals or groups, but it suggests that this elite group maintained a continuous relationship with the Eurasian steppe.

### Genetic data on the social structure of the Avar society

In this study, we produced novel information regarding the social organisation of the Avar elite stratum. Considering the variability of the maternal and paternal lineages, the Y chromosomal profiles give a contrasting picture to the mitochondrial ones. We gained Y-STR information on seven out of eight males of the Kunbábony group. They all belong to the Y haplogroup N-Tat, but at least to three different N-Tat haplotypes. One N-Tat haplotype is shared not only among the highest elite males, but is also found in other six elite males, all buried at Kunszállás. All individuals that have this identical Y-STR profile were buried in the Danube-Tisza Interfluve (Fig. 1). One other N-Tat lineage is limited to a single site (Kunpeszér), but connects all investigated individuals on that burial ground. Based on our data this Avar-period elite group shows strong biological ties, possible paternal kinship relations. Therefore, we conclude that the Avar elite probably inherited their power and wealth through the paternal line. Paternal kinship was also an organizing rule within the communities of two studied sites, Kunszállás and Kunpeszér.

The Avar society has been understood in the framework of nomadic societies^2^. It is widely accepted that kinship ties (both real and fictitious) were of higher importance for nomads than for settled groups. Kinship is a social segment that is defined based on the proximity of individuals to each other in the system of biological relationship. Among the nomadic societies of Central Asia strict patrilineality has been observed, but matrilineal lineages were also recorded and noted in certain cases. Kinship is also a way of understanding the world and creating order in it, and also served as the framework within the social order was maintained^38–40.^

While nomadic societies are described as segmentary, they are not necessarily egalitarian. In segmented societies the rank and the relationship between individuals and/or groups is determined as well as their place in the wider society, thus a system is created, where no one has his/her exact equal. The kinship system differentiates between superior and inferior lineages and emergence of a dominant lineage could occur. The paternal kinship relations among the investigated individuals buried in lavishly furnished graves indicate the presence of such a dominant lineage in the Avar society. The importance of kinship in nomadic societies has been challenged, but never in the case of the elite strata^41^. The idea of a chosen or sacred segment is a known political notion in the Central Asian nomadic societies^34,38–40^.

Based on our current knowledge about the previous populations of the Carpathian Basin, we presume that the Asian components of the Avar elite entered the region with the Avar conquest. Considering that the investigated Avar elite group was at least 3-4 generations younger than the time of the Avar conquest, mitochondrial DNA of both males and females gives us valuable information about social structure of the Avar period elite.

Our results suggest that the Avar elite did not mix with the local 6^th^ century population for ca. a century and could have remained a consciously maintained closed stratum of the society.

The dominance of the Asian mtDNA lineages (especially in males) suggest, that only after that period did the number of intermarriages with local women increase, and the Avar elite was mostly endogamous (within the strata or Avars of Asian origin) in the Carpathian Basin. Moreover, while it does not contradict the models of elite migrations, it shows that the Avar elite arrived in family groups, or at least men and women migrated together.

It is important to note that the investigated elite group consists of mostly male burials (n=18); the women belonging to the same social strata are archaeologically barely visible. From the investigated sites in the Danube-Tisza Interfluve, only one female individual was buried with high value artefacts; the other richly furnished female burials are located in the Transtisza region. The male power appeared in the public sphere, while the female power manifested probably in the family sphere. This did not mean however, that women could not wield public power/influence^40,42^, but could have led to different representational forms. To get more insights, and define the uniqueness of the Avar period elite’s paternal and probably maternal gene pool, we need to study the common people of the Avar society as well.

## Conclusion

We present here the first complete mitogenome and Y-STR dataset from the Avar period of the Carpathian Basin. Our results attest that the maternal and paternal genetic lineages of the Avar period elite in the Carpathian Basin was different from the European uniparental genepool of their period, and was mostly of East-Central Asian origin. The detected East-Central-Asian maternal and paternal genetic composition of the elite was preserved through several generations after the Avar conquest of the Carpathian Basin.

This result suggests a consciously maintained closed society, probably through internal marriages or intensive contacts with their regions of origin. The results also hold valuable information regarding the social organisation of the Avar period elite. The mitochondrial DNA data suggests that not only a military retinue consisting of males migrated, but an endogamous group of families. The Y-STR information support that the Avar elite was organized by paternal kinship relations, and kinship had also an important role in the usage of the elite’s cemeteries. The kinship relations among the investigated elite individuals buried in lavishly furnished graves indicate the presence of a dominant lineage that correlates to the known political notion of chosen or sacred segment of nomadic societies.

Our first genetic results on the leader class of the Avar society provide new evidence of the history of an important early medieval empire. Nevertheless, further genetic data from ancient and modern Asian populations and from the Avar period of the Carpathian Basin is needed to describe the genetic relations and the genetic substructure of the Avar-period population in greater detail.

## Materials

The studied individuals were excavated at ten different sites (found in small burial groups or single burials). Seven out of ten sites are located in the Danube-Tisza Interfluve^7,14,15^. The primary focus of the sample selection was to target all available members of the highest elite group of the Avar society.

Out of the 26 investigated Avar period samples, eight individuals show similar archaeological characteristics with weaponry covered with precious metal foils, ornamented belt sets and drinking vessels made of gold or silver (Csepel-AC1, Kecskemét-AC23, Kunbábony-AC2, Kunpeszér Grave 3-AC21, 8-AC22, 9-AC20, Petőfiszállás Grave 1-AC19, Szalkszentmárton-AC8). The wealth of the 50-60 years old male from Kunbábony is outstanding with the 2.34 kilograms of gold buried with him (Fig. 2, SI chapter 1a).

During the sample collection it became evident that these individuals are also tied together by their physical anthropological characteristics, as the skulls showed certain morphological traits, that are not characteristic to the 6^th^ century local populations and are rare in the 7^th^ century as well^43,44^(SI chapter 1d). To have a better understanding of this group, we later collected samples from 18 individuals from the same region and the neighbouring Transtisza region with similar morphological ancestry skull remarks (see Table S1), but without any outstanding grave goods unique only to the highest social ranks (Fig. 1, Fig. S11).

## Methods

### Ancient DNA work

Twenty-six samples were collected from ten different cemeteries dated to the Avar period (7^th^-8^th^ centuries) according to their geographical position, grave goods, funerary custom and anthropological characteristics (see Table S1 and the site and grave descriptions in the Supplementary Information).

All stages of the work were performed under sterile conditions in a dedicated ancient DNA laboratory (Laboratory of Archaeogenetics in the Institute of Archaeology, Research Centre for the Humanities, Hungarian Academy of Sciences) following well-established ancient DNA workflow protocols^12,45^. The laboratory work was carried out wearing clean overalls, facemasks and face-shields, gloves and over-shoes. All appliances, containers and work areas were cleaned with DNA-ExitusPlus™ (AppliChem) and/or bleach and irradiated with UV-C light. All steps were carried out in separate rooms. In order to detect possible contamination by exogenous DNA, one extraction and library blank were used as a negative control for every batch of five/seven samples. Haplotypes of all persons involved in the ancient DNA work were determined and compared with the results obtained from the ancient bone samples.

Usually, pars petrous bone fragments were used for analyses, except for three individuals where teeth and long bone fragments were collected because the skulls were not preserved (Table S1).

The DNA extraction was performed based on the protocol of Dabney et al.^46^ with some modifications described also by Lipson et al.^45^. DNA libraries were prepared using UDG-half treatment methods^47^. We included library negative controls and/or extraction negative controls in every batch. Unique P5 and P7 adapter combinations were used for every library^47,48^. Barcode adaptor-ligated libraries were then amplified with TwistAmp Basic (Twist DX Ltd), purified with AMPure XP beads (Agilent) and checked on a 3% agarose gel. The DNA concentration of each library was measured on a Qubit 2.0 fluorometer. In solution, the hybridisation method was used to capture the target short sequences that covered the whole mitochondrial genome, as described by Haak et al. and Lipson et al.^8,45^. Captured samples as well as raw libraries for shotgun sequencing were indexed using universal iP5 and unique iP7 indexes^48^. NGS sequencing was performed on an Illumina MiSeq System using the Illumina MiSeq Reagent Kit v3 (150-cycles).

AmpFLSTR Yfiler PCR Amplification Kit (Thermo Fisher Scientific) was used for the Y chromosome STR analyses. We followed the instructions of the manufacturer’s user manual, except applying 34 cycles for PCR amplification instead of the standard 30 cycles protocol. Fragment analyses of PCR products were performed on a 3130 Genetic Analyzer in accordance with the manufacturer’s instruction. Data evaluation, allele sizing and genotyping were determined by using GeneMapper® ID v3.2.1 software (Applied Biosystems). We amplified and analysed each sample at least twice and alleles were designated according to the parallel analyses with minimum detection threshold at 50 RFU.

STR results are seen in Table S1. Y haplogroups were predicted using www.nevgen.org and the predicted terminal SNPs were checked on the Y tree of ISOGG version 14.04 (https://isogg.org/tree/). We searched for haplotype matches in YHRD database (YHRD.ORG by Sascha Willuweit & Lutz Roewer).

### Bioinformatics analyses

The final BAM files were obtained by a custom pipeline for both shotgun and capture datasets. The paired-end reads were merged using SeqPrep master (https://github.com/jstjohn/SeqPrep), allowing a minimum overlap of 5 bp and minimum length of 15 bp. Then, the reads were filtered by size and barcode content using *cutadapt* version 1.9.1^49^, allowing no barcode mismatch, and a minimum length of 15 bp. BWA version 0.7.12-r1039^50^ was used to map the capture sequencing reads to the Cambridge Reference Sequence (rCRS) and the shotgun sequencing reads to hg38 human genome assembly allowing a 3 bp difference in seed sequence, 3 bp gap extension and 2 gap opening per reads. The downstream analyses including SAM-BAM conversion, sorting, indexing and PCR duplicate removal was performed by *samtools* version 1.6^51^.

For capture data, indel realignment was performed using Picard tools version 2.5.0 (https://github.com/broadinstitute/picard) and GATK version 3.6^52^. The presence of a deamination pattern was estimated by MapDamage version 2.0.8 (https://ginolhac.github.io/mapDamage/) and summarised in Table S1. Due to the relatively young age and half-UDG treatment of the samples required, the deamination frequency did not reach the minimum limit for software *schmutzi* in most cases; therefore, the final validation of the sequences was performed by eye on the final bam files. The shotgun sequencing provided a raw estimate of the endogenous content and genetic sex determination according to Haak et al.^8^. These data are summarised in Table S1.

The consensus sequences (with a minimum coverage of 3x) and SNP calls according to rCRS^53^ and RSRS^54^ (with a minimum variant frequency of 0.7 and minimum coverage of 5x) were generated by Geneious 8.1.7 software (https://www.geneious.com/). The whole mitochondrial fasta sequences were submitted to the NCBI GenBank. The haplogroups were determined using HaploGrep (v2.1.1) (https://haplogrep.uibk.ac.at/) based on Phylotree version 17^55^.

### Population genetic analyses

Standard statistical methods were used for the calculation of genetic distances between the investigated Avar elite population and Eurasian ancient and modern populations.

We excluded sample AC20 from any statistical and phylogenetic analyses because of the large number of haplogroup-diagnostic positions missing. Furthermore, we excluded sample HC9 from population-genetic statistical analyses because it belongs to a later period (end of 7^th^ – early 9^th^ centuries), and also excluded sample RC26 from sequence-based analyses because of a large number of unreadable and missing parts of the mitochondrial sequence, which inhibit the haplotype-based calculation of genetic distances.

The whole mitochondrial genomes of the samples were aligned in SeaView^56^ by *ClustalO* with default options. Positions with poor alignment quality were discarded in the case of ancient and modern sequences as well.

Population pairwise F_ST_ values were calculated based on 4,015 modern-day and 1,096 ancient whole mitochondrial sequences using Arlequin 3.5.2.2^57^. The Tamura and Nei substitution model was used^58^ with a gamma value of 0.62, 10,000 permutations and significance level of 0.05 in case of comparison between the investigated Avar elite population and 43 modern-day Eurasian populations (for the references see Table S5). For the comparison of 26 ancient populations, the Tamura and Nei model was performed with a gamma value of 0.599, 10,000 permutations and significance level of 0.05. The number of usable loci for distance computation in this case was 13,526 because 3,021 np had too much missing data (for the references see Table S4). The genetic distances of linearised Slatkin F_ST_ values^59^ were used for Multidimensional scaling (MDS) and visualised on two-dimensional plots (Figs. 3-4) using the metaMDS function based on Euclidean distances implemented in the vegan library of R 3.4.1^60^.

Principal component analyses (PCA) were performed based on mtDNA haplogroup frequencies of 64 modern and 48 ancient populations. Thirty-two mitochondrial haplogroups were considered in the PCA of ancient populations, while 36 mitochondrial haplogroups in the PCA of modern populations were considered (Tables S2-S3). The PCAs were carried out using the prcomp function in R 3.4.1 and visualised in two-dimensional plots displaying the first two (PC1 and PC2) or the first and third principal components (Figs. S4a-b and S5a-b).

For hierarchical clustering, Ward type algorithm^61^ and Euclidean distance measurement method were used based on haplogroup frequencies of ancient and modern populations and displayed as a dendrogram in R3.4.1 (Figs. 2 and S6). The same population-pools were used for this clustering as those used in the two PCAs.

### Phylogenetic analysis

Phylogenetic analyses aimed to detect close maternal relative lineages within the group of samples belonging to a certain haplogroup. All available human mitochondrial genome sequences in NCBI (more than 33,500) were downloaded and sorted according to their haplogroup assignments. Multiple sequence alignment was performed for each sample set using the same procedure mentioned in the Population genetic analyses section, with an exception that only the 303-318 sites were discarded on this highly repetitive and indel prone region due to poor alignment quality. Then neighbour joining trees were calculated using the *dnadist* and *neighbor* subprograms of Phylip version 3.696^62^ with default options. The Median Joining Network, which is a favoured method for analysing haplotype data, was rejected due to unresolvable ties. The trees were drawn in Figtree version 1.4.2 (http://tree.bio.ed.ac.uk/software/figtree). We did not use bootstrap analyses due to the low quantity of informative positions, which highly biases the supporting values.

To examine the Y-STR variation within the Y chromosomal haplogroups, Median Joining (MJ) networks were constructed using the Network 5.0 software (http://www.fluxus-engineering.com). Haplogroups predicted as 162 N-Tat samples from 12 populations, 127 N-F4205 samples from six populations and Q1b-M346 samples from 15 populations were included in the networks (see Table S9-11 for references and for the Y-STR used in the analyses). Post processing MP calculation was applied, creating network containing all shortest tree. Repeats of the locus DYS389I were subtracted from the locus DYS389II and, as is common practice, the locus DYS385 was excluded from the network. Within the network program, the rho statistic was used to estimate the time to the most recent common ancestor (TMRCA) of haplotypes within the compared haplogroups. Evolutionary time estimates were calculated according to Zhivotovsky et al.^63^ and STR mutation rate was assumed to be 6.9×10-4 /locus/25 years.

## Acknowledgements

This study was funded by a Hungarian research grant of T.V. (NKFIH-OTKA NN 113157) based on ELTE – Eötvös Loránd University, Budapest. We thank further support to the Szilágyi Family Foundation. ASN was supported in her work by the János Bolyai Research Fellowship of the HAS. We thank Ildikó Pap, Ágnes Kustár (Hungarian Natural History Museum, Budapest), the co-workers of the Tessedik Samuel Museum (Szarvas) and the Katona József Museum (Kecskemét) for providing DNA samples. We are grateful to Csilla Balogh (Istanbul Medeniyet University, Istambul), and Erika Wiker (Katona József Museum, Kecskemét) for providing archaeological description and information about the studied sites. We thank Levente Samu and Viktor Szinyei for preparing the base maps. We are grateful to Walter Pohl, Patric Geary, Csanád Bálint and Jan Bemmann for reading and commenting the manuscript.

## Author contributions

I. K, B.G.M, A.S-N, T.V. designed the study. V.Cs., G.D, B. Sz. and B. E. performed the ancient DNA analyses. V.Cs. G.D., H. P. and A.S-.N, performed population genetic analyses. I.K., G.Cs., A.G., B.K., G.M.L., G.L. and T.V. performed the archaeological evaluation, provided the historical background and interpretation. A.M, B.G.M., E. M. and Gy.P. summarised the anthropological data of the human remains. V.Cs., D.G., I.K, G.Cs., H. P., B.G.M, A. S-N, T.V wrote the paper. All authors read and discussed the manuscript.

