## Supplementary text and figures for "Genetic insights into the social organisation of the Avar period elite in the 7^th^ century AD Carpathian Basin"

#### Supplementary Information

#### Index

### 1. Supplementary text

#### a) Description of the Avar period archaeological sites

(by A. Gulyás, B. Kovacsóczy, G. M. Lezsák, G. Lőrinczy, T. Vida and I. Koncz)

**Békésszentandrás –Benda-tanya, site 76. (Békés county, Hungary)**
(by A. Gulyás, G. Lőrinczy)

Two Avar-age burials were excavated at Békésszentandrás (46°52'14.06"N; 20°28'50.18"E) during the motorway construction in 2015. An elderly female[a] was found in the lavishly furnished niche Grave 87 which was oriented NW-SE. She was lying in a coffin gouged out of a tree-trunk, although the grave was disturbed, it was still richly furnished.

To the end of the side niche a grey, good quality jug turned on a fast wheel was placed. Under the coffin and inside the niche elements of a horse harness were found: mounts, bits and stirrups. The particularity of Grave 87 from Békésszentandrás is that the horse harness was adorned with 84 gold and copper four-lobed mounts and 11 strap-ends with interlace pattern. Based on their parallels from Fönlak, Törökkanizsa and Tiszavasvár-Kashalom the pressed mounts date the burial to the middle of the 7th century.

The grave structure suggests that the deceased buried at Békésszentandrás were descendants of groups arriving from the Eastern European steppes to the South part of the Transtisza region during the Avar conquest of the Carpathian Basin. These groups buried skinned, but harnessed horses together with certain individuals. The deposition of horse harness elements without the animal only appeared later, at the middle of the 7th century and became common at the end of the century. The musculoskeletal stress markers on the bones of the upper arm and thighs suggest that the female individual used to ride regularly, so it is no coincidence that the whole set of horse harness was placed beside her into the grave.

The goat sacrum and the lumbar vertebrae found in Grave 24 and 25 from Szarvas and Grave 87 from Békésszentandrás were placed into the burials as food offerings. In the Transtisza region during the early Avar period animal bones (or meat) as food offerings are very rare, and almost without exception sheep or goat sacrum, caudal or lumbal vertebrae. This tradition survives in the cemeteries of the Körös-Tisza-Maros Interfluve until the second half of the Avar period.

The unpublished archaeological and anthropological material are stored in the Tessedik Sámuel Museum, Szarvas.

---

[a] The physical anthropological analysis was conducted by Antónia Marcsik.

**Budapest –Csepel –Kavicsbánya**[1]
(by I. Koncz, T. Vida)

The lavishly furnished burial of a 45-50 years old male[2] was found during mining works in 1924 on the Csepel Island (47°16'33.17"N; 18°57'29.76"E). Only parts of the funerary assemblage made it to the museum. He was buried with a sword decorated with gold plates. The P-shaped suspension loops were decorated with inlays framed by pearl-rows. The decoration of the sword and its accessories has a very close analogy in Grave 8 from Kunpeszér, the only difference is that on the inner side of the suspension loops—instead of one—two inlays are present. A small bronze buckle was probably part of the sword suspension as well. Three 3-bladed arrowheads and one 2-bladed arrowhead also came to light. Iron staples prove the presence of a coffin.

The high-ranking individual was probably a member of the Avar period elite belonging to the military retinue of the *Qagan* and as such settled in the vicinity of the *Qagan*'s seat.

The archaeological material is stored in the Budapest History Museum, Budapest and the anthropological material in the Department of Anthropology, Hungarian Natural History Museum, Budapest.

**Kecskemét – Sallai utca (Bács-Kiskun county, Hungary)**[3]
(by T. Vida)

The burial was found during canalization works in 1973 in the inner city of Kecskemét (46°54'46.44"N; 19°42'17.19"E). The adult male[3] was buried with his ring-pommel sword decorated with gold sheets and gold-foiled silver rosettes. At the outer side of his left arm the remains of a quiver were found with ten pieces of three-bladed arrowheads. Around the pelvis area elements of a garnished belt were excavated: gold-plated silver mounts (shield-shaped, double shield-shaped and crescent-shaped mounts) and a strap-end. The belt belongs to the so called Pančova-type and is dated to the middle third of the 7$^{th}$ century. The closing part of his pouch hanging from his belt was a reused suspension of a Byzantine type lamp.

The archaeological material is stored in the Katona József Museum, Kecskemét while the anthropological material in the Department of Anthropology at the University of Szeged, Szeged.

**Kunbábony (Bács-Kiskun county, Hungary)**[4]
(by T. Vida)

The burial was found and destroyed during sand mining works in 1971. As a result, *in situ* observations were not possible. The archaeological and anthropological material was recollected from the workers on the following days. Despite the disturbance, the ’princely’ burial from Kunbábony (47° 1'5.85"N; 19°13'16.62"E) still has the richest burial assemblage from the early Avar period with its 155 preserved artefacts. It contained 2.34 Kg of gold, the second being Bócsa with its 1.33 Kg.

From the Kunbábony burial the remains of a senile (approx. 60 years old) male with Mongoloid characteristics came to light[5]. He had an injury just under his left eye that healed but rendered him half blind for the rest of his life. He was buried into a coffin decorated with gold sheets; the coffin was placed on a bed. The exact size of the NW-SE oriented burial is unknown.

An important element of the Avar period male representation is the use of double belt sets (a weapon belt and one with decorative and representative function only). One of his belts consisted of pseudo buckles with glass inlays joined with an imported Byzantine buckle made of gold with mosaic-like decoration. His other belt was adorned with gold mounts decorated with granulation (Fig. 2). The man was buried with his ring-pommel sword, dagger, bow and 25 arrowheads. His weapons – except the arrowheads – were decorated with gold sheets and half-palmette motifs. A birdhead-shaped gold sheet could be interpreted as part of a sceptre or horsewhip. A drinking horn—its parallels are known from Mala Pereshchepina (Bulgaria) and from the treasure of Nagyszentmiklós (Sânnicolau Mare, Romania)—and a gold jug represent the tableware. An amphora of Byzantine origin – the volume of which is 50 l – was placed into the burial as drink offering, while the gold-plated wooden bowls together with sheep bones suggest the presence of food offerings. The crescent-shaped gold sheet—probably placed on the chest as a pectoral—shows direct Central-Asian connections.

The burial rite, the funerary attire and the grave goods point to the East as elements of the steppe nomadic material culture of the period. The technological details and the decoration however suggest a culturally heterogeneous origin. The grave goods from the burial represent a high-ranking individual from the Eurasian steppe who got attracted by the Byzantine-Mediterranean world as the Byzantine elements are only superficially present embedded into nomadic traditions. The gold pseudo buckles with pearl-row decoration were made with early Byzantine techniques, probably by Byzantine workshops or goldsmith working in the Carpathian Basin and were popular among the Avar and other Eastern-European elite. The amphora and the Byzantine buckle with mosaic-like decoration suggest gift exchange as part of a diplomatic relationship.

The dating of the Kunbábony burial is based on the relative chronology of the grave goods. Certain ornamental characteristics present on the belt mounts (chip carving, pearl rows, the shape of the stone inlays etc.) with direct parallels from Bócsa and Petőfiszállás suggest an earlier dating to the first half of the 7th century.

The burial from Kunbábony is the most prominent of the group of elite burials from the early Avar period located in the Danube-Tisza Interfluve. The high rank of the deceased is indicated by his wealth and personal objects, but because of the lack of a unequivocal *regalia* compared to the already mentioned Mala Pereshchepina, Kunbábony cannot be interpreted as the *Qagan* known from the historical sources.
The archaeological and anthropological material is stored in the Katona József Museum, Kecskemét.

**Kunpeszér – Felsőpeszéri út (Bács-Kiskun county, Hungary)**[6]
(by T. Vida)

During the construction of a road in 1894 a pyramid-shaped gold earring was found. On the same site (47° 4'39.11"N; 19°14'33.45"E) a rescue excavation was conducted between 1982 and 1984 and 27 graves were excavated in two groups. 15 burials (6 males, 6 females and 3 children) belong to the first group; they have N-S orientation and are laying remarkably far (20-40 meters) from each other. Most of the males and females buried in the richly furnished graves had Mongoloid characteristics[7].

Five males were buried with gold-plated swords with P-shaped suspension loops and bows. The suspension loop of the sword from Grave 3 is the closest parallel to the one known from the high-ranking burial at Csepel. Their belts were decorated with mounts and strap-ends made of silver. The rite of Charon's obol and the presence of goat/sheep bones as food offerings were also observed in the cemetery. Jewellery made of precious metals (earrings with spherical pendants, pyramid-shaped earrings etc.) and objects of everyday use (mirrors, pouches, capsules etc.) are characteristic for female burials.

The first burial group of Kunpeszér belongs to the Kunbábony – Bócsa group. This small cemetery dated to the middle of the 7$^{th}$ century can be interpreted as a group of high-ranking families belonging to the military retinue of the *Qagan*.

The second burial group contained 16 disturbed burials with individuals showing Europid morphological traits. These poorly furnished (or even without any kind of grave goods) burials probably belonged to the lower social strata of the Avar society. This cemetery group is dated to the beginning of the 8$^{th}$ century and its connection with the first group is still unclear.

The archaeological material is stored in the Katona József Museum, Kecskemét while the anthropological material in the Department of Anthropology at the University of Szeged, Szeged.

**The Avar-period cemetery of Kunszállás-Fülöpjakab (Bács-Kiskun county, Hungary)**[8]
(by G. M. Lezsák)

The cemetery of Kunszállás-Fülöpjakab (46°45'2.92"N; 19°44'16.68"E) was excavated by Elvira H. Tóth between 1967 and 1979. The cemetery lies on a NW-SE oriented sand hill covering an area of 900 $m^2$. The 61 graves with NW-SE orientation were organised into loose rows. 13 males, 22 females, 5 juveniles, 20 children - were buried at the site[9,10]. Based on the grave finds the cemetery was in use from the second half of the 7th until the end of the 8th century.

The cemetery is well represented in male burials with garnished belts. The chronological frame of the site is given based on the technical details, form and decoration (pressed, plated-cast with griffin and tendril pattern). The burials were also richly furnished with weapons: mainly bows, sabres, swords and knives came to light.

The higher social status of certain individuals is represented by artefacts made of gold. The men from Grave 32 was buried with gold earrings, golden hair clips, gold-foiled strap-ends with interlace pattern made of silver and a sword decorated with gold plates (second half of the 7th century). The burial with its prestigious grave goods could be rated to the single burials of the military elite from Kecskemét-Miklóstelep, Ballószög, Szeged-Átokháza (the so-called Ozora-Tótipuszta group), at the same time its belt set also connects it to the earlier Kunbábony-Bócsa group. From Grave 52 a gold-foiled cast bronze belt set with griffin and floral ornament came to light. The inwrought bronze belt mounts with gold-foiling and palmette motif prove that the cemetery survived until the second half of the 8th century. Compared to the small number of the burials at the site, the percentage of belt sets pressed of gold sheets (Graves 24 and 32) and gold-foiled bronze belts (Graves 20, 52 and 59) are very high.

The cemetery could be interpreted as the burial site of members of the military retinue and their families in the close vicinity of the political centre. Cemeteries in the region (Hetényegyháza, Tatárszentgyörgy, Kunadacs, Kunbaracs, Kecskemét-Ballószög, Szabadszállás-Batthyány utca, Gátér, Kiskőrös-Cebe, Kiskőrös-Vágóhíd, Tiszakécske-Óbög, illetve a Homokmégy-halom, Solt-szőlőhegy, Madaras-téglavető, Szeged-Fehértó B.) suggest that this area remained a seat of power until the 8th century.

The archaeological analysis of the cemetery is in progress by Gabriella Lezsák (HAS RCH MŐT), the anthropological analysis is also in progress by Antónia Marcsik.

**Petőfiszállás (Bács-Kiskun county, Hungary)**[11,12]
(by I. Koncz, T. Vida)

The richly furnished male burial at Petőfiszállás (46°37'15.34"N; 19°50'37.74"E) was found by metal-detecting connected to the construction of the M7 motorway in 1998. Part of the burial was disturbed during its discovery. The 40-45 years old male had Mongoloid characteristics. The area around the burial was later extensively studied, but no more graves were found.

The high-ranking male was buried with a sword covered with gold plates and had two belt sets. One of the belts consisted of shield-shaped, lunula-shaped, oval- and T-shaped mounts and two strap-ends made of gold with pearl-row decoration. The belt widens on the back marked with a tripartite composite mount. This belt set is a close parallel to the belt sets of the Bócsa-Kunbábony group in both decoration and technology, but the lack of the so-called pseudo buckles suggests an earlier dating. This composition appears in the second fourth of the 7th century, but it is possible that its use overlapped with the use of the pseudo buckles during the middle of the 7th century. Based on the analysis of Éva Garam the gold belt set of Petőfiszállás belongs to the Bócsa-Kunbábony group, but predates other sites with pseudo buckles such as Mala Pereshchepina and Maglód. The other belt set was decorated with round, partially gilded silver mounts, that has the closest parallels in Bócsa.

The importance of the burial from Petőfiszállás comes from its belt sets that connect the early horizon of Avar period belts (abstract floral and point-comma ornaments etc.) with the Bócsa-Kunbábony group characterized by pearl-row decoration and pseudo buckles.

The burial belongs to the group of elite burials with military character located in the Danube-Tisza Interfluve that could be interpreted as the military retinue of the *Qagan*.

The archaeological material is stored in the Katona József Museum, Kecskemét, the anthropological material is stored in the Department of Anthropology at the University of Szeged, Szeged.

**Szalkszentmárton – Táborállás Grave 1. (Bács-Kiskun county, Hungary)**
(by B. Kovacsóczy)

Since October of 2015 a complex, multi-layered site is being excavated at Szalkszentmárton (county Bács-Kiskun) by the Katona József Museum of Kecskemét. Settlement traces from the Neolithic, bronze-age cremation burials and two Avar period graves were found in the outskirts of the town at Táborállás (46°58'33.34"N; 19°01'01.62").

Grave 1 is a N/NW-S/SE oriented pit burial (l: 227 cm; w: 151 cm; d: 90 cm) with a niche on its west side. An adult (23-39 years old) male with Mongoloid characteristics came to the light from the burial.[b] The skin, skull and lower legs of his harnessed horse were also buried with him.

The burial from Szalkszentmárton is the first from the Kunbábony group that was excavated by archaeologists and documented *in situ*. Both the custom of burying a horse and the niche structure were unknown from this group before.

The man was buried with two belt sets. One of them consisted of round belt mounts made of bronze with a lead core; their middle part was gold-plated decorated with circle-dot ornament. These mounts have parallels in the Bócsa-Kunbábony-Petőfiszállás group. The other belt consisted of small shield-shaped mounts, double crescent-shaped mounts and small strap-ends with mask pattern. These mounts show direct contact to the Martynovka type belt sets. Parallels of the ’T’-shaped mounts are known from Kunbábony and Bócsa. The deceased was buried with a ring-pommel sword, probably with a bow (as the badly preserved bone plates suggest) and five arrowheads. Based on the grave goods, the burial could be dated to the first half/mid-7$^{th}$ century.

Szalkszentmárton lies at a strategically important position: it was a Danube crossing since the Bronze Age and during the Roman and Late Roman period (*Intercisa*) as well. The high-ranking man from Grave 1 was probably responsible for the control and protection of the river crossing.

The unpublished archaeological and anthropological material is stored in the Katona József Museum, Kecskemét.

**Szarvas –Kovács-halom site 8/1. (Békés county, Hungary)**
(by, A. Gulyás, G. Lőrinczy)

During the construction of an irrigation system a row of six W-E oriented Avar period burials was excavated at Szarvas–Kovács-halom in 2015 (46°48'4.74"N; 20°35'6.63"E). Four of the simple pit graves contained noteworthy grave goods. The two adult females (Grave 25 and 33) a mature male (Grave 24) and a juvenile, possibly female individual (Grave 40) all belonged to the Mongoloid taxonomic group based on their morphological traits[c].

Three graves contained the very same elements of the horse harness: curb bits and iron stirrups with straight foot. The females from Grave 25 and 40 also had a harness decorated with mounts. In Grave 25 pressed, four-loped mounts and tufted strap-ends decorated with pressed-in rhombus lines were found. From Grave 40 pressed rosette mounts and rare mounts with tulip-like shape came to the light that only has contemporary parallels from the cemetery

[b] The physical anthropological analysis was conducted by Antónia Marcsik.
[c] The physical anthropological analysis was conducted by Antónia Marcsik.

of Zamárdi. The man from Grave 24 was not furnished with decorated harness, but possessed a belt with rectangle shaped pressed mounts. Based on the grave goods the burials could be dated to the second half of the 7th century.

Round gold sheets were placed inside the mouth of the woman (Grave 25) and the girl (Grave 40) as a replacement of Charon's obol. The need for the replacement of coins appeared after the expiry of inflow of Byzantine coins into the Carpathian Basin during the second half of the 7th century.

The goat sacrum and the lumbar vertebrae found in Grave 24 and 25 from Szarvas and Grave 87 from Békésszentandrás were placed into the burials as food offerings. In the Transtisza region during the early Avar period bones (or meat) as food offerings are very rare, and almost without exception sheep or goat sacrum, caudal or lumbal vertebrae are found. This tradition survives in the cemeteries of the Körös-Tisza-Maros Interfluve until the second half of the Avar period.

The unpublished archaeological and anthropological material are stored in the Tessedik Sámuel Museum, Szarvas.

**Székkutas-Kápolnadűlő (Csongrád county, Hungary)**[13]
(by T. Vida)

Katalin B. Nagy conducted a rescue excavation between 1965 and 1986 at the Kápolnadűlő area of Székkutas (46°30'21.12"N; 20°32'16.39"E) uncovering around 18,000 $m^2$. A cemetery of 535 graves dated to the Avar period (7th-9th centuries) came to light, that is till today the only entirely excavated cemetery of the Avar period in the Middle-Tisza region. The site gives a comprehensive picture about the material culture of the Avar period in the Transtisza region as it was used continuously from the 7th until the 9th century.

The senile male was laid to rest in a WNW-ESE oriented niche burial (Grave 396)[14]. The niche was built at the shorter side of the grave. The elements of the garnished belt were found between the bones. The belt was adorned with a profiled bronze buckle, an iron strap-holder, rectangular mounts with curved sides made of bronze, a bronze strap-end and a complex backplate consisting of five mounts. The middle mount of the backplate was decorated with gold-foiling on the sides and with a blue stone inlay in the middle. It was complemented by two shield-shaped on its longer sides and two triangular mounts with traces of gold-foiling on the shorter sides. Remains of a wooden vessel were also observed in the burial.

The Grave 396 is dated to the end of the 7th century based on the belt set. Although the cemetery was founded in the middle Avar age, many early Avar period characteristics are recognizable: niche burials, animal sacrifices, burials with horse harnesses, the presence of goat/sheep sacrum or lumbar vertebrae etc.). Grave forms and the burial rite changed drastically in the 8th century. In that sense Grave 396 showcases a transitional period between the early Avar and middle Avar ages.

The archaeological material is stored in the Móra Ferenc Museum in Szeged while the anthropological material in the Department of Anthropology at the University of Szeged.

#### b) East-Central Asia as a geographic framework

Central Asia was used as a general term for the eastern part of the continental mass of Eurasia. East-Central Asia is applied to the eastern part of Central Asia and it largely corresponds with the term Inner Asia as used by Nicola Di Cosmo[15]. In geographical terms, it includes three areas: Manchuria (China) in the east; Mongolia in the centre, including parts of Gansu, northern Shanxi, and northern Shaanxi; and today's Xinjiang in China, the Minusinsk Basin and the northern Altai Mountains in the west (Russia).

#### c) Archaeological and historical records of the origin and migration of the Avars, and a centre of power in the Danube-Tisza Interfluve

(by T. Vida, G. Csiky, I. Koncz)

The Avars united the peoples of the Carpathian Basin under their rule at the end of the 6th century. The historical sources suggest that this group was bringing titles and institutions of a nomadic state to the Carpathian Basin. The written account of Theophylact Simocatta describes a Byzantine embassy to the Western Turkic ruler, Silzibulos. According to his claim the Avars were Turkic subjects who escaped from his will and that they have unrightfully usurped the title Qagan and the name 'Avar' as these fugitives were not real Avars in the Turkic Qagan's view[16].

As the predecessor of the Turkic Qaganate was the Rouran Empire in the East-Central Asian steppes, the idea of Rouran origin of the Avars appeared as early as the 18th century[16,17]. A rival hypothesis is based on the same source mentioning the alternate name of the migrating group as Warkhonitai which would connect them to the Hephtalites of Hunnic origin from Central Asia[18]. Recent historical research understands these groups not as homogeneous ethnicities, but rather as entities connected by common political and economic goals.

The concentration of burials in the Danube-Tisza Interfluve containing weapons, such as ring-pommel swords decorated with gold sheets and bows and a rich array of prestige items including belt sets and tableware made of precious metals can in all likelihood be linked to a leading group of the early Avar polity and maybe even to the Qagan's military retinue (Kunbábony-Bócsa group). The single burial of a 50-60-year-old man at Kunbábony (AC2) contained 2.34 kilograms of gold. Because of its outstanding wealth it is often described as the 'burial of the Qagan'; while it cannot be clearly proved archaeologically (eg. there are no royal/Qagan's insignia and diplomatic gifts in the grave), he was indeed a prominent member of the society. The artefacts found in these burials point to both eastern (steppe) and Mediterranean cultural traditions, suggesting that the group had far-reaching connection networks and were incorporating elements of different origin to its representation[4,19].

The crescent-shaped gold sheet from the Kunbábony burial have direct analogies from 5th-6th century burials attributed to the Rouran from Mongolia and to the Northern Wei dynasty (Tuoba Xianbei) in Northern China where these artefacts were used as pectorals based on their documented position in burials (Yihe Nur in China[20] and Talyn Gurvan Kherem in Mongolia[21]), although these items were regarded earlier as head-gears or diadems used as insignia[22] (Fig. S3).

Characteristic ostentatious edged weapons covered with golden or silver sheets were found in early Avar elite burials, and were probably used as prestige goods. These ring-pommel swords have good contemporaneous analogies from the Altai Region, but their distribution reached as far as China, Korea and Japan, while their representations can be found in mural paintings of Old-Samarkand (Afrasiab)[23,24] (Fig. S1).

The strong East-Central Asian connection of the early Avar period material culture is present on the level of the common people as well (see the rectangular-mouthed vessels and vessel with peaked and knobbed rim[25]). The technological breakthrough of the iron stirrup, already known in Asia, appeared in Europe at the same time as the Avars; its importance was also showed by the cavalry reform of the Byzantine military after its large-scale distribution[26–28]. However, not only certain artefact types reached Europe during that time, but also some characteristic ways of deposition also spread: sacrificial assemblages so-called ‘tainiks’ (cache) composed of weapons and horse harnesses were buried in shallow pits in the Carpathian Basin during this period[23,29–31] (Fig. S2).

#### d) Anthropological characteristics of the Avar period population in the Carpathian Basin

(by B. G. Mende, A. Marcsik, E. Molnár and Gy. Pálfi)

The Avar period is both well represented and researched from a physical anthropological point of view. The 250-270 years long period has around 100,000 excavated burials, from which about 10,000 were analysed anthropologically. The main reason behind the outstanding anthropological interest is that the so called Asian craniometric and morphologic types appear in large quantities and in high diversity for the first time in the Carpathian Basin. Their typology, spatial and temporal distribution became one of the defining components of the anthropological picture of the Avar population, that has had an effect on the historical-archaeological narratives as well.

The Avar period population of the Carpathian Basin was anthropologically heterogeneous, mainly characterized by European (previously Europid/Caucasoid in scientific literature) craniometric and morphologic types with both brachycranic and dolichocranic variations. Important parts of this heterogeneity are the Asiatic types—so called Mongolid types in the Soviet-Russian and East-European tradition —as well. The main observation of Pál Lipták's work from 1983[32], that Asiatic types did not exceed 15-20% of the Avar period population—even if in certain areas/sites it could go up as far as 50%—regarding the Danube and Tisza Interfluve, is valid still today[33,34]. The craniofacial morphologic traits characteristic to Asiatic skull types—shovel-shaped incisors, the typical "rocking" form of the mandible, the presence of mandibular torus and palatine torus, the shape of the nasal bones, the width of the supranasal region—could appear on other types on occasion as well[35–37]. The main distinction between the European and Asiatic skull types is based on the morphological and metric characteristics of the upper part of the facial bones and the forehead. These kind of analyses have been used by both Soviet-Russian and Eastern-European anthropologists –hallmarked by the work of Debec and Alekseev[38] – to distinguish populations with Asian origin.

Skull types connected to Asian populations occur in almost every early Avar period cemetery. While the appearance of these Asiatic types in the early Avar period can be interpreted through the arrival of new, possibly East-Central Asian population groups, their presence in the middle and late Avar periods is still intensively debated.

#### e) Detailed results of the mitochondrial DNA phylogenetic analyses

(by V. Csáky, D. Gerber, A. Szécsényi-Nagy)

In the followings, general and phylogenetic information is summarised on each mtDNA haplogroup that was detected in the Avar dataset.

Haplogroups C and D, detected in the Avar elite in a frequency of 33%, are the most common haplogroups throughout Northern, Central and Eastern Asia. The oldest lineages expanded before Last Glacial Maximum from East-Asia, and from there their dissemination indicates the post-glacial recolonization of North-Asia. Both haplogroups were involved in migration from East-Asia and southern Siberia to East- and Northeast Europe probably during the middle Holocene[39]. Phylogenetic tree of C4a1a (Fig. S10a) contains three samples from the Avar period (AC7, AC13 and AC15 from cemetery Kunszállás) of which AC13 and AC15 have basal position to all other 20 East-Asian C4a1a4 individuals. The C4b tree (Fig. S10b) has a C4b6 branch containing an Avar sample AC1 from Csepel cemetery and further modern Central and East-Central Asian individuals (Uyghurs, Buryat, two Altaian Turkic individuals such as an Altai-Kizhi and a Tubalar). The trees of haplogroups D (D4i, D4j and D5b, Figs. 7, S10c-d) contain five samples from cemeteries Kunbábony (AC2), Békésszentandrás (AC3), Szarvas-Kovácshalom (AC4 and AC6), and Kunpeszér (RC26), and show the Central- and North Asian connection through clustering of the Avar mitogenomes mostly with modern Uyghurs living in Northwest China, Yakuts, Buryats, Chinese and individuals from the Pamir region (Table S6). On the D4i tree, the Avar individual AC6 is in basal position of a diverged branch that contains East-Central Asian and North Asian present-day individuals (Fig. 7).

The haplogroup F is most common in East-Asia and Southeast-Asia, while subclade F1b is frequent in Central Asia too (empop.org). Phylogenetic tree of F1b1 haplogroup signals genetic connection of the Avars with Eastern-Siberians and Inner-Asians (Fig. S10e): samples AC11 and AC12 from Kunszállás neighbouring with two Chinese individuals and AC21 clusters with similar haplotypes of Uyghur and Tungus individuals, the latter living in East-Siberia.

Haplogroup M7 is characteristic especially for East Asian populations[40], whereas sub-haplogroup M7c1b2b has been detected among Chinese, Uyghurs, Malays and Thai[41–44] (Table S6). The closest relative of AC22 is a Southeast Asian individual, and the structure of the phylogenetic tree suggests a Southeast Asian origin of the sub-haplogroup as well. However, due to the low number of available samples, the existence of another unsampled or already extinct more north-western branch of M7 cannot be rejected (Fig. S10f).

Haplogroup R2 is detected from Near East to South-East-Asia: in Iran, Georgia, Pakistan, China, Thailand and in Central-South Asian populations (Makrani, Balochi, Brahmani and Pamirs)[45,46]. Among these populations, the sample AC18 from Kunpeszér clustered together again with an Uyghur individual from Northwest China on the phylogenetic tree of R2 (Fig. S10g).

Haplogroup Y is detected in higher frequency in populations Nivkhi, Ulchi, and Hezhen (Far Eastern Russia) and Nias (Indonesia)[46,47]. The phylogenetic tree of haplogroup Y1a indicates basal position of sample AC23 from Kecskemét-Sallai út compared to Y1a1 samples from other Asian populations, namely Evenks, Uyghurs, Yakuts, Buryats, Chinese, and also Hezhen and Udegey living in the Far Eastern Russia. This tree topology signalizes a common ancestor of this maternal lineage in these populations (Fig. S10h).

Haplogroup H is the most frequent haplogroup in present-day Europe, it encompasses ca. 34-40% of the mitochondrial variability[48]. H is also common in North Africa and the Middle East. We detect two sub-haplogroups of H: the phylogenetic tree of H5a signals an entirely European character for AC5 from Szarvas-Kovácshalom (Fig. S10i), while the H8 tree represents strong Caucasian connection of AC17 from Kunpeszér (Fig. S10j).

The J and T lineages expanded from Near East to Europe at the time of the Neolithization[49,50]. The phylogenetic trees with two investigated samples (AC8 from Szalkszentmárton and HC9 from Székkutas) belonging to these haplogroups (J1b and T1a1b) that signal typical European character too (Figs. S10k and S10l). We did not construct a phylogenetic tree for haplogroup T1a1 because too many modern individuals with different origin belong to this haplogroup and as such, it seems to be phylogeographical mixed and not resolvable.

U5 is one of the oldest European haplogroup, which is detected in European Late Glacial and Holocene hunter-gatherers in the highest frequency[51]. Two subgroups appear among the Avars: U5a1 (sample AC16 from Kunpeszér) (Fig. S10m) and U5b1b (AC10 from Kunszállás), of which phylogenetic trees indicate dispersed Eurasian phylogeographic pattern (Fig. S10n).

Haplogroup Z1a is widespread in North-Europe, Volga-Region, as well as in Siberia[52]. The here presented phylogenetic tree of Z1a haplogroup contains interesting divergence of Asian and North-European lineages, where the investigated individual AC19 from cemetery Petőfiszállás is basal to the Asian modern samples (Fig. S10o). It might indicate that AC19 shared common ancestor with maternal ancestors of present-day Uyghur, Evenk, Yakut, Yukaghir and Nganasan individuals.

#### f) Detailed results of the Y-STR network analyses

(by H. Pamjav and A. Szécsényi-Nagy)

Based on ten Y-STR loci, networks were constructed within N-Tat (N1a1-M46 or N1c or earlier named as N3) and N-F4205 (N1a1a1a1a3a-F4205) haplogroups. Ten STRs overlapped among the populations studied. All included haplotype and haplogroup results can be found in Table S9.

An MJ network of 162 N-Tat haplotypes is shown in Fig. 3. The founder haplotype I (Cluster 1) is shared by eight populations including three Mongolian, three Székely, three Northern Mansi, two Southern Mansi, two Hungarian, eight Khanty, one Finnish and two Avar (AC17, AC26) chromosomes, thus Cluster 1 included 24 samples. Haplotype II (Cluster 2 in Fig. 3) include 45 haplotypes from six populations studied: 32 Buryats, two Mongolians, one Székely, one Uzbek, one Uzbek Madjar, two Northern Mansi and six Avars (AC15, AC12, AC14, AC1, AC19 and KSZ 37). Cluster 1 (allele 11 in DYS391) differs one molecular step from Cluster 2 (allele 10 in DYS391). We consider Cluster 1 to be the founder lineage because this haplotype is shared by eight populations and more diverse than the Cluster 2. Derenko et al. first revealed this observation in Siberian populations[53]. One Avar haplotype (AC8; see red arrow in Fig. 3) seems to be a descendent from other haplotype cluster that is shared by three populations including six haplotypes (two Buryat, three Khanty and one northern Mansi samples). This haplotype cluster also differs one molecular step (locus DYS393) from haplotype II. Most Hungarian N-Tat samples are located on the top of the network together with samples originated Ural Region belonging to different subgroup. Most Buryat males share the same haplotype (Fig. 3, Cluster 2), which phenomenon is due to genetic drift and indicates haplotype II to be younger than haplotype I. All N-Tat samples included in the network were tested for N- L708 SNP and showed derived allele for L708 SNP as well.

Based on our calculation, the age of accumulated STR variance (TMRCA) within N-Tat lineage for all samples is 7.0 kya (thousand years ago) (95% CI: 4.9 - 9.2 kya), considering the core haplotype (Cluster 1) to be the founding lineage. This estimated TMRCA is much younger than its sequence based calculation of 13 kya (95% CI: 11.3-14.6 kya)[54]. However, the time estimate for the N-L708 subclade is 8.4 kya in Ilumäe et al.[54]. We note that all our N-Tat samples tested by SNP M46 were also derived for SNP L708 supporting the observation of Ilumäe et al. (2016) and justifies our TMRCA calculation (7.0 kya), which approaches the age of the L708 (8.4 kya).

The TMRCA for only Avar and Hungarian N-Tat samples is 3.5 kya (95% CI: 1.4 - 5.7 kya). It is interesting that the lower limit of the confidence intervals fits to the time when the Avars lived in Carpathian Basin (600 AD). If we also calculated the TMRCA for the Avar and Buryat samples, it was 4.6 kya (95% CI: 1.8 – 7.4 kya), which is little bit older than that of the Avars and Hungarians.

Then, we aimed to classify the Avar samples to downstream subgroup within the N-Tat haplogroup and again constructed a network including N-F4205 (N1a1a1a1a3a-F4205) samples tested by Ilumäe et al. (Fig. S4, Table S10)[54]. According to the Y-chromosomal haplogroup tree of Ilumäe et al. study, three sub-clades can be distinguished under N-Tat (named as N3a-L708, N3b-B187 and N3c-B496 in the paper), for us the only N3a-L708 (N3a= N1a1a1a on the ISOGG tree version 14.04) branch can be interesting. The authors identified further bifurcations under N-L708 (N1a1a1a-L708) haplogroup. From them, the N-F4205 (earlier N3a5-F4205) is mostly present in the Mongolian, Buryat and Kazakh populations; thus, we considered that our Avar samples probably belong to this subgroup. Our network results support this assumption (Fig. S4, Cluster 1 and 2), because eight out of nine Avar samples share the same haplotypes with the N-F4205 samples tested by us and Ilumäe et al.[54] on modern samples.

Based on our calculation, the age of accumulated STR variance (TMRCA) within N-F4205 lineage for all samples is 6.3 kya (95% CI: 3.2 - 9.3 kya), considering the core haplotype (Cluster 1) to be the founding lineage which is older than that of Ilumäe et al.[54]. The TMRCA for only the Avar and Buryat N-F4205 samples is 4.6 kya (95% CI: 1.3 - 7.9 kya), which is absolutely in agreement with the 4.6 kya calculated by Ilumäe et al.[54]. The lower limit of the confidence intervals fits to era when the Avars lived in the Carpathian Basin (ca. 700 AD). As in the case of the N-Tat samples, we also calculated the age for Hungarians and Avars and it was 4.4 kya (95% CI: 1.6-7.2 kya).

As seen in Fig. S4 (Cluster 1 and 2), the presence of common haplotypes of the Hungarian speakers and Avars may indicate a link to physical connection in their history. There can be two possibilities for this observation: first, some Hungarian N-F4205 samples may be descendants of the Avars as it is fact that Avars lived in Carpathian Basin before arriving the Hungarian Conquerors. The second, the ancestors of these Hungarian males could come from Asia with ancient Hungarians later at the end of the 9$^{th}$ century AD. At present, there is no scientific method to show such time differences.

The frequency peak of N-F4205 (N3a5-F4205) chromosomes is in Transbaikal region of Southern Siberia and in Mongolia[54]. Thus, we conclude that most Avar N-Tat (F4205) chromosomes probably originated from this area and completely in line with the results of NGS Y-chromosomal sequences published by Ilumäe et al.[54] According to them, the split of N3a5-B197 (upstream SNP of the F4205) into sub-lineages occurred soon after the emergence of the clade itself (nearly 5.0 kya), distributing N3a5-F4205 within Mongolic-speaking populations and N3a5-B202 among populations of the Chuckotko-Kamchatka language family[54], which is consistent with our results. In addition, the Avar samples shared common haplotypes with Hungarian speakers within N-Tat (L708) and N-F4205 subhaplogroups supporting a genetic link between some of recent Hungarian speakers (especially from Hajdú-Bihar county and Hungarian speaking Székely males in Romania) and the Avar people.

At the same time, the reliable historical and genetic conclusions require an extension of the study to a significantly larger database with deep haplogroup resolution, including more ancient DNA data and NGS sequencing of Y chromosomes.

#### 2. Supplementary Figures

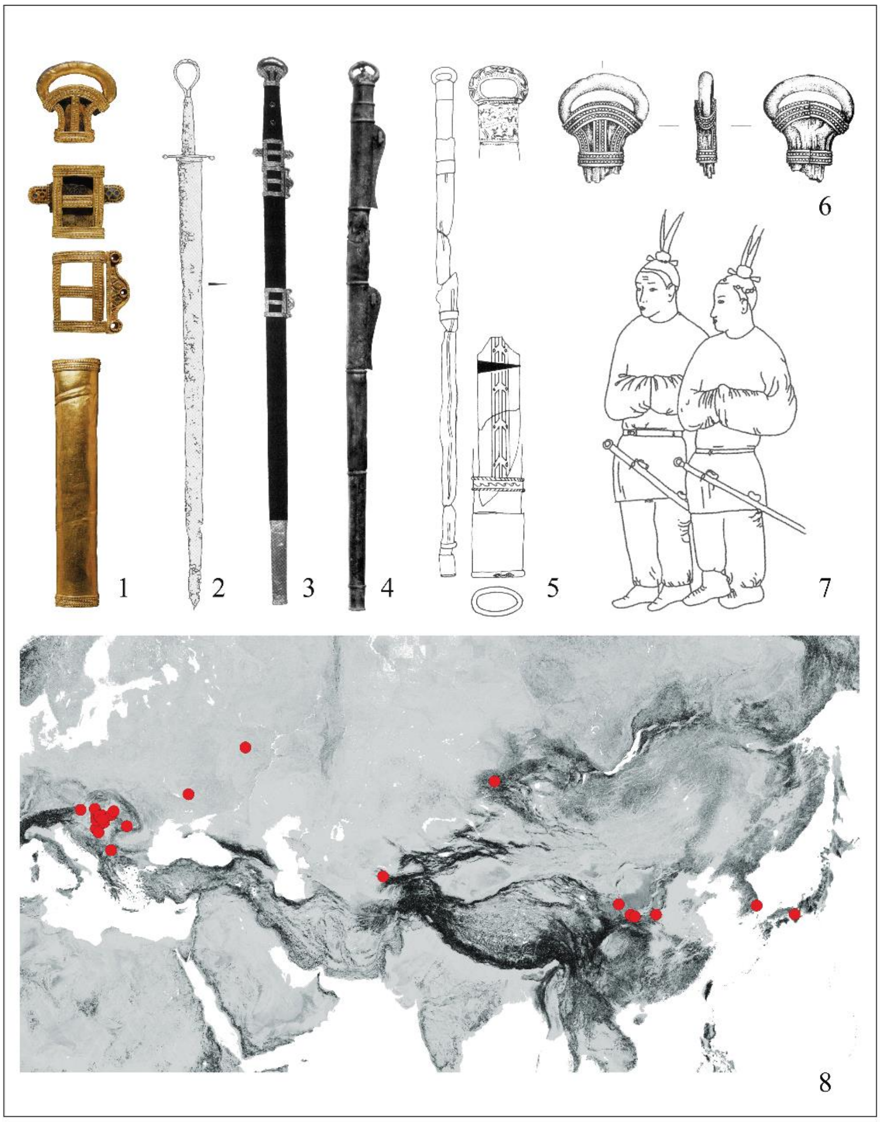

**Figure S1. Ring-pommel swords.**

Kunbábony[4], 2. Sopron-téglagyári agyagbánya[24], 3. Bócsa[24], 4. Miniature sword from the burial of Emperor Wu of Northern Zhou, Xian-yang, Xi'an, Shaanxi province[55], 5. Gyeongju, South Korea, grave No. 126[56] 6. Bócsa[24], 7. Afrasiab 1. building: representation of Korean envoys (after Al'baum 1975[57] and Csiky 2015[24]), 8. Distribution of the ring-pommel swords[24]. Pictures and drawings were taken from the cited literature. Background map is based on the Global Multi-resolution Terrain Elevation Data 2010 (GMTED2010), available from the U.S. Geological Survey (https://lta.cr.usgs.gov/GMTED2010) without restrictions to use.

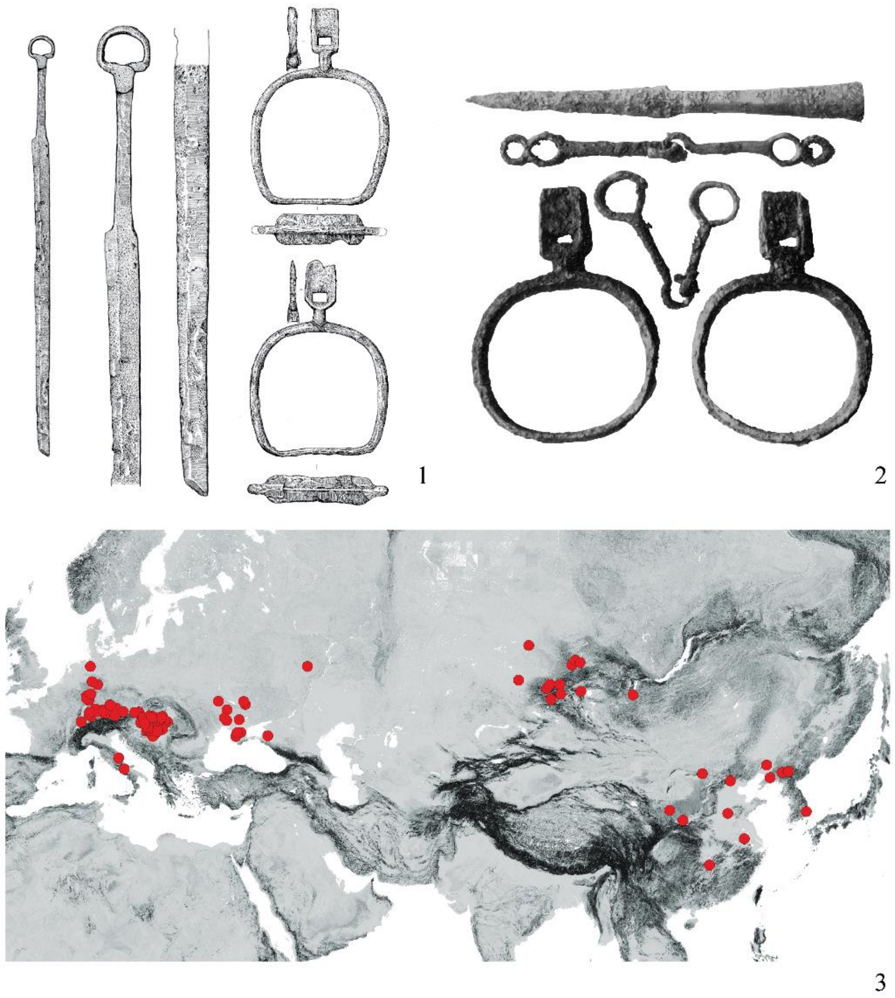

**Figure S2. Sacrificial assemblages and stirrups.**
Ak Kaja, Altai Republic, Russia[23]; 2. Csengele-Jójárt, Hungary[58]; 3. Distribution of iron stirrups between the $4^{th}$-$7^{th}$ centuries (after[59], changed by G. Csiky and I. Koncz). Pictures and drawings were taken from the cited literature. Background map is based on the Global Multi-resolution Terrain Elevation Data 2010 (GMTED2010), available from the U.S. Geological Survey (https://lta.cr.usgs.gov/GMTED2010) without restrictions to use.

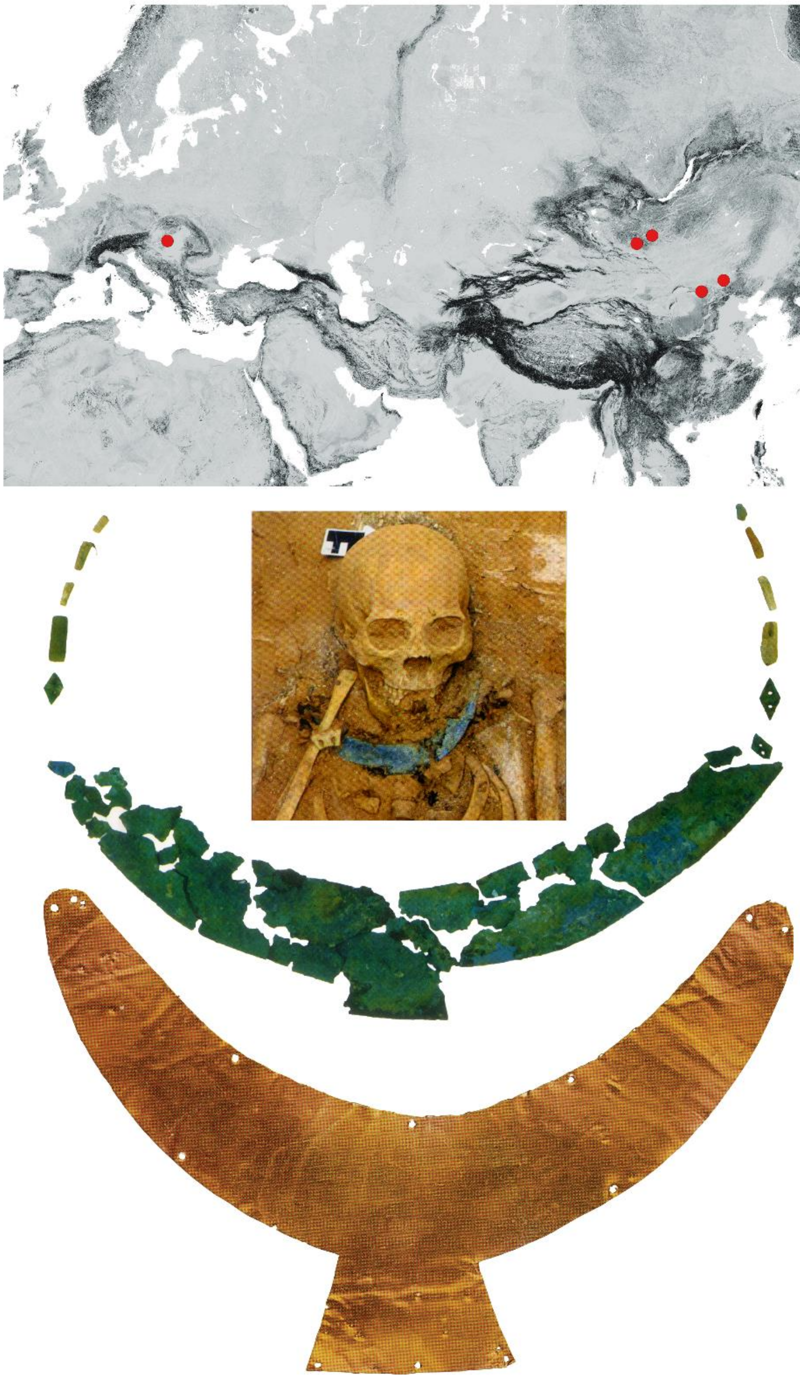

1

3

2

**Figure S3. Crescent-shaped gold sheets used as pectorals in graves of the nomadic elite.** Distribution of the crescent-shaped gold sheets used as pectorals from East to West: Yihe Nur[20], Bikeqi[22]; Talyn Gurvan Kherem (with photos in figure part 3)[21], Galuut sum[22], Kunbábony (pictured in figure part 2)[4]. Pictures were taken from the cited literature. Background map is based on the Global Multi-resolution Terrain Elevation Data 2010 (GMTED2010), available from the U.S. Geological Survey (https://lta.cr.usgs.gov/GMTED2010) without restrictions to use.

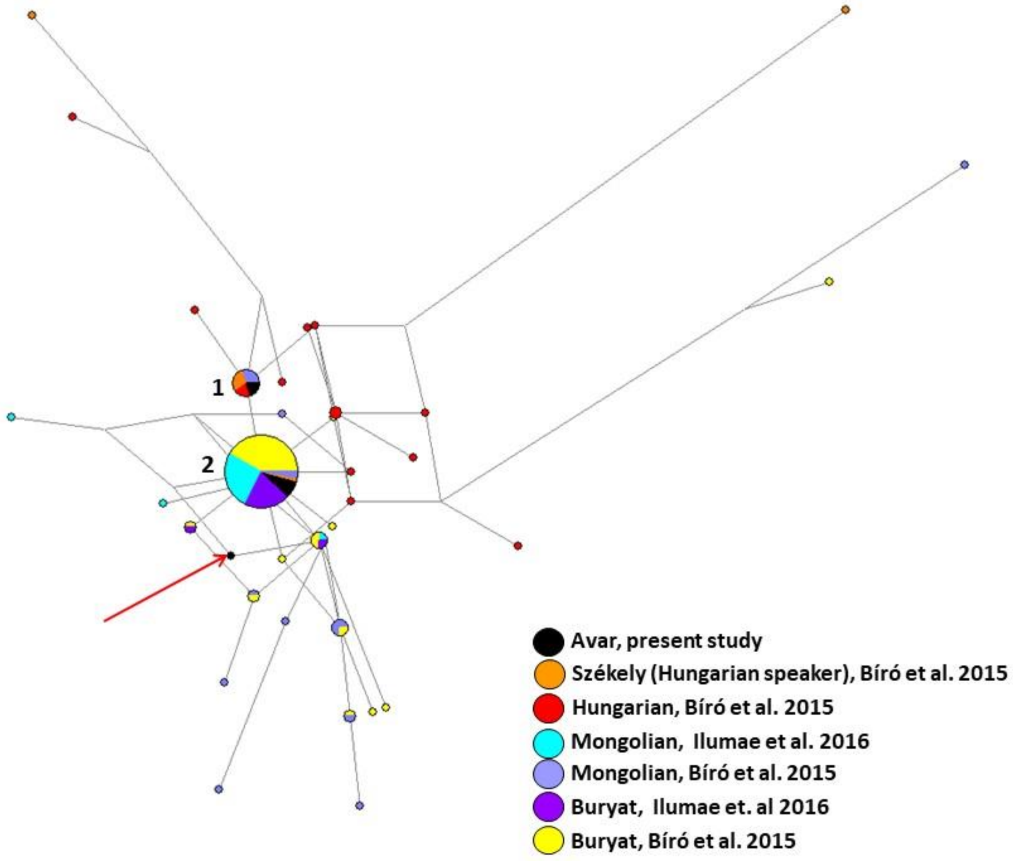

**Figure S4. MJ network including N-F4205 Y-STR haplotypes.** Cluster 1 includes two Avars (AC17 and AC26) together with three Mongolian, three Székely and two Hungarian males. Cluster 2 includes 74 haplotypes belonging to the following populations: 15 Mongolian and 19 Buryat N-F4205 haplotypes[54], 31 Buryat, two Mongolian, one Székely[60] and six Avar (AC15, AC12, KSZ37, AC14, AC1 and AC19) samples of the present study. Remaining one Avar sample (AC8; highlighted by a red arrow) is two mutational steps away from Cluster 2. Most modern Hungarian samples can be derived from Cluster 1 by one or more mutational events. Information on the modern samples is seen in Table S10.

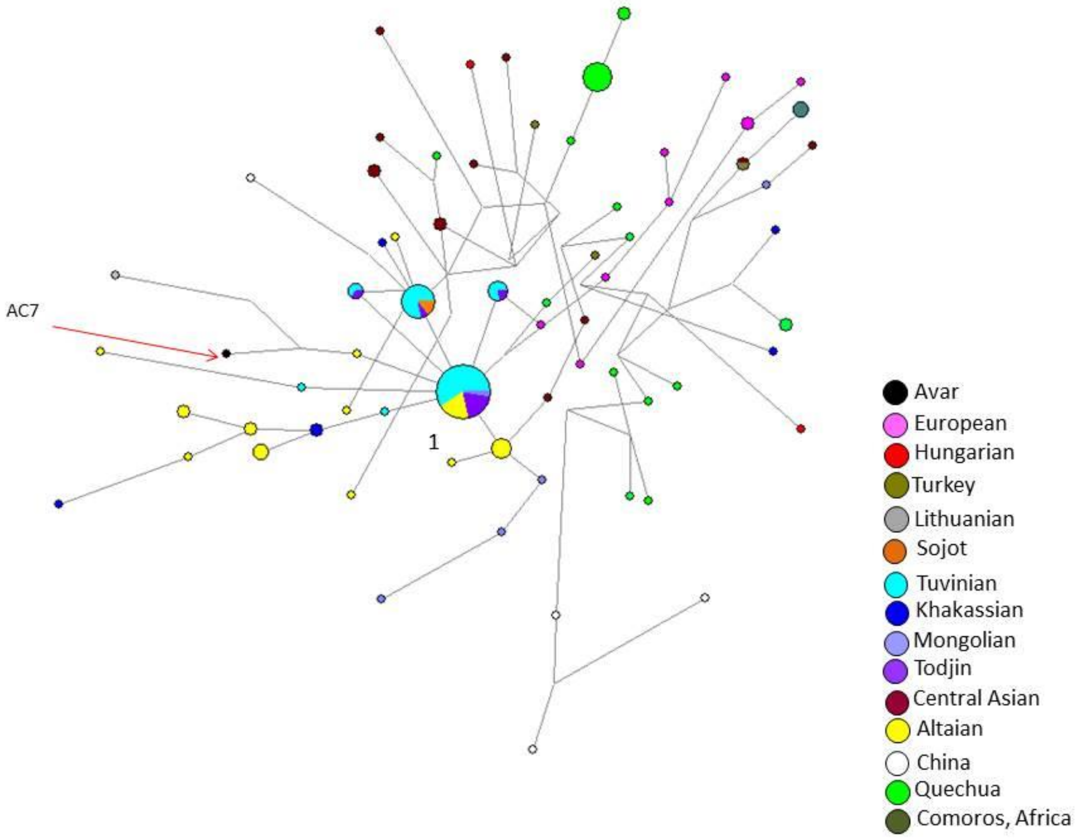

**Figure S5. MJ network including Q1b-M346 Y-STR haplotypes.** One Avar sample (AC7; highlighted by a red arrow) is on one branch with an Altaian individual and a Lithuanian. In Cluster 1 seven Altaian, seven Todjin, 22 Tuvinian and one Mongolian individual can be found. The AC7 is in a distance of three STR mutations (DYS389, DYS392, DYS389) from this cluster, which branch is led through a Tuvinian male. References are listed in Table S11.

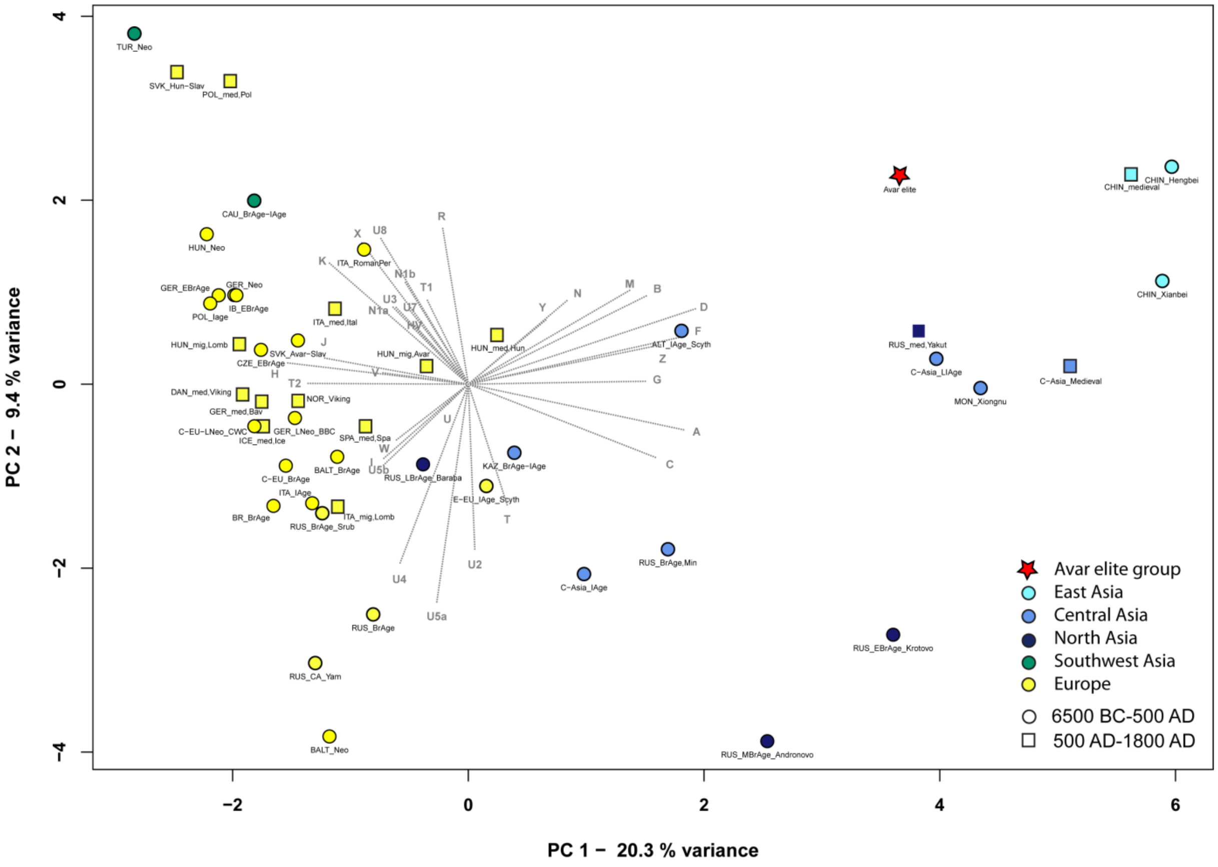
PC 2 – 9.4 % variance
PC 1 – 20.3 % variance
4
2
0
−2
−4
−2
0
2
4
6
Avar elite group
East Asia
Central Asia
North Asia
Southwest Asia
Europe
6500 BC-500 AD
500 AD-1800 AD

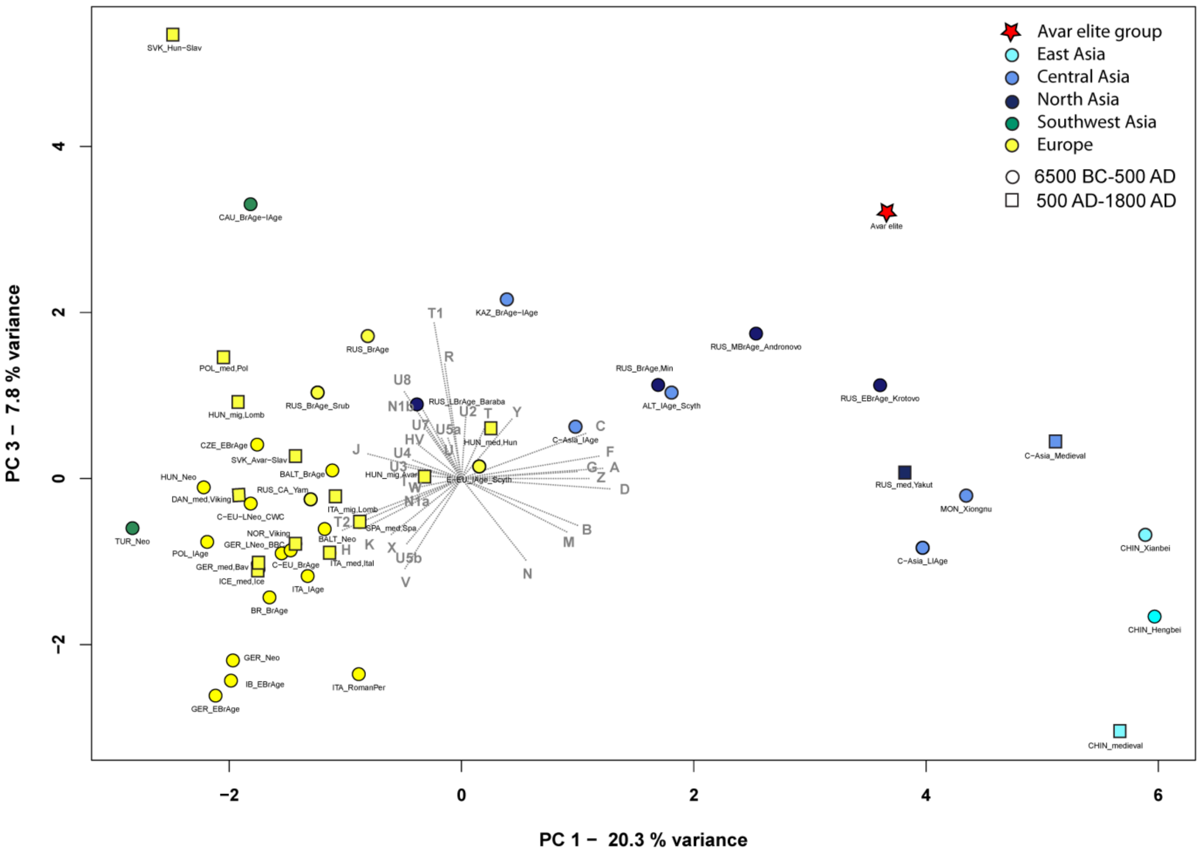
PC 3 – 7.8 % variance
PC 1 – 20.3 % variance
4
2
0
−2
−2
0
2
4
6
Avar elite group
East Asia
Central Asia
North Asia
Southwest Asia
Europe
6500 BC-500 AD
500 AD-1800 AD

**Figure S6. PCA plots with 48 ancient populations, representing first two principal components (A) and first-third principal components (B); (37.5 % of the total variance is shown).**

The differentiation of European and Asian populations is displayed on the PCA plot of ancient populations along PC1, PC2 and PC3. The separation is caused by the opposite influence of Asian (A, B, C, D, G, F, Z) and European (H, J, K, T2) haplogroups along PC1, while R and the subgroups of haplogroup U (especially the U4, U5a and U8) predominate on PC2. The Avar elite group is situated close to 5$^{th}$ – 3$^{rd}$ centuries BC Scythians from Altai region (ALT_Scythians), medieval Chinese population and Yakuts from 15-19$^{th}$ centuries AD (RUS_Yakuts) along PC1 and PC2, while along the PC3 Avars are near to Siberian populations of Andronovo (SIB_Andronovo) and Krotovo (SIB_Krotovo) cultures in Asian part of plot. For the haplogroup frequencies, abbreviations and references see Table S2.

A

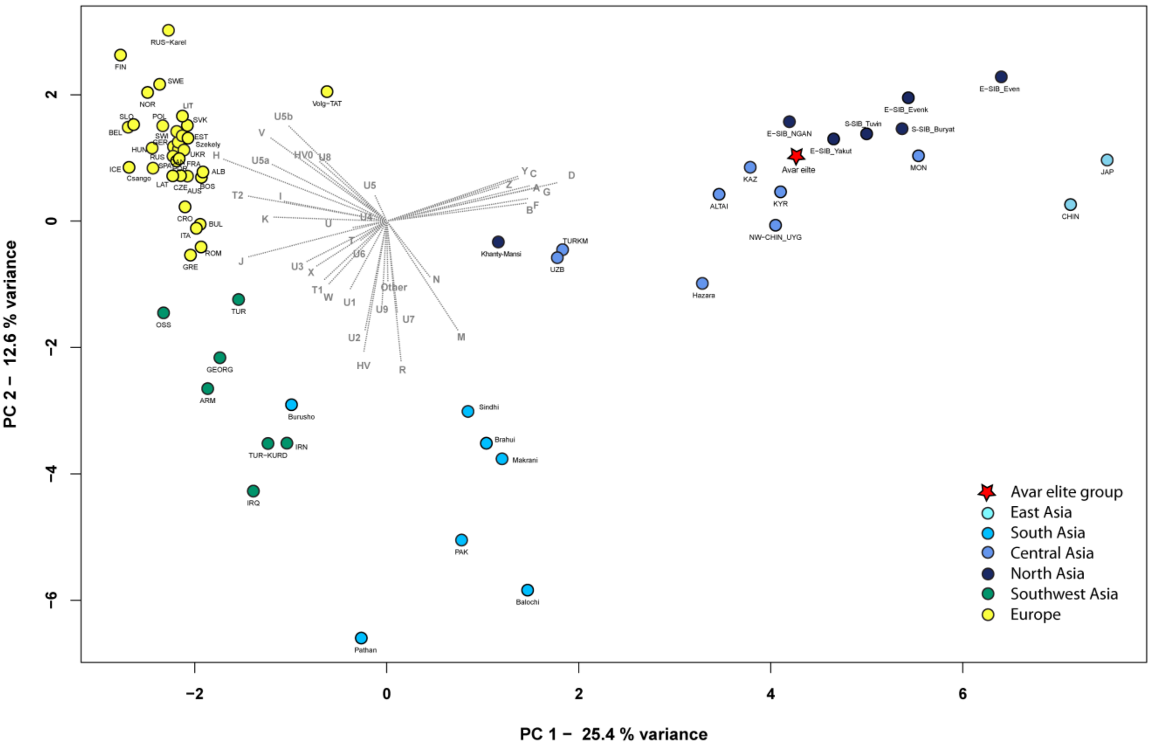

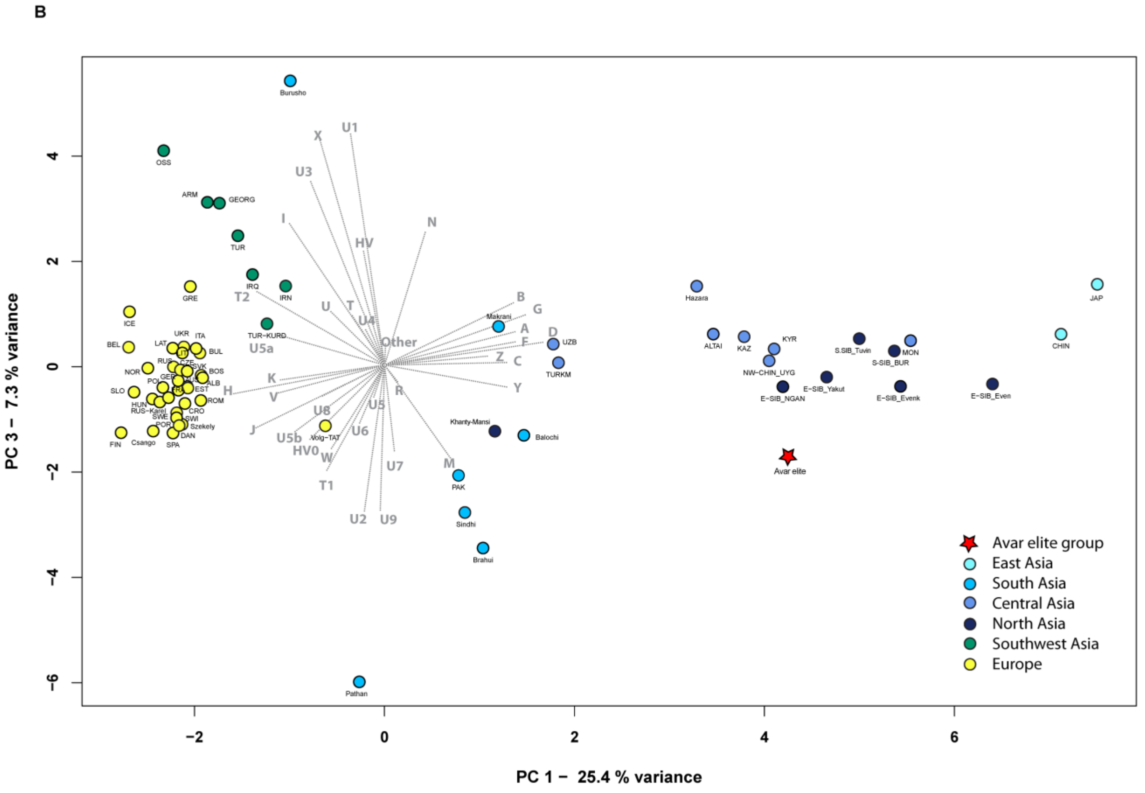

**Figure S7. PCA plots of the first two principal components (A) and first-third principal components (B) comparing haplogroup frequencies of 64 modern populations; (45.3 % of total variance is shown).**

The PCA plots shows separation of European, Asian, Near-Eastern and some Central-Asian populations. The European populations cluster along PC1 and PC2. Southwest Asian populations and populations of South-Central Asia are separated along PC2 from Europe, but PC3 differentiates Southwest Asia and South-Central Asia from each other. The Asian populations are differentiated and dispersed along PC1. Along these three components the Avar elite group is clustered together with Central Asian populations from Kazakhstan (KAZ), Kyrgyzstan (KYR) and Altai region (ALTAI) as well as with Uyghurs living in Northwest-China (NW-CHIN_UYG) and East-Siberian Yakuts (E-SIB_YAKUT) and Nganasans (E-SIB_NGAN). For the haplogroup frequencies, abbreviations and references see Table S3.

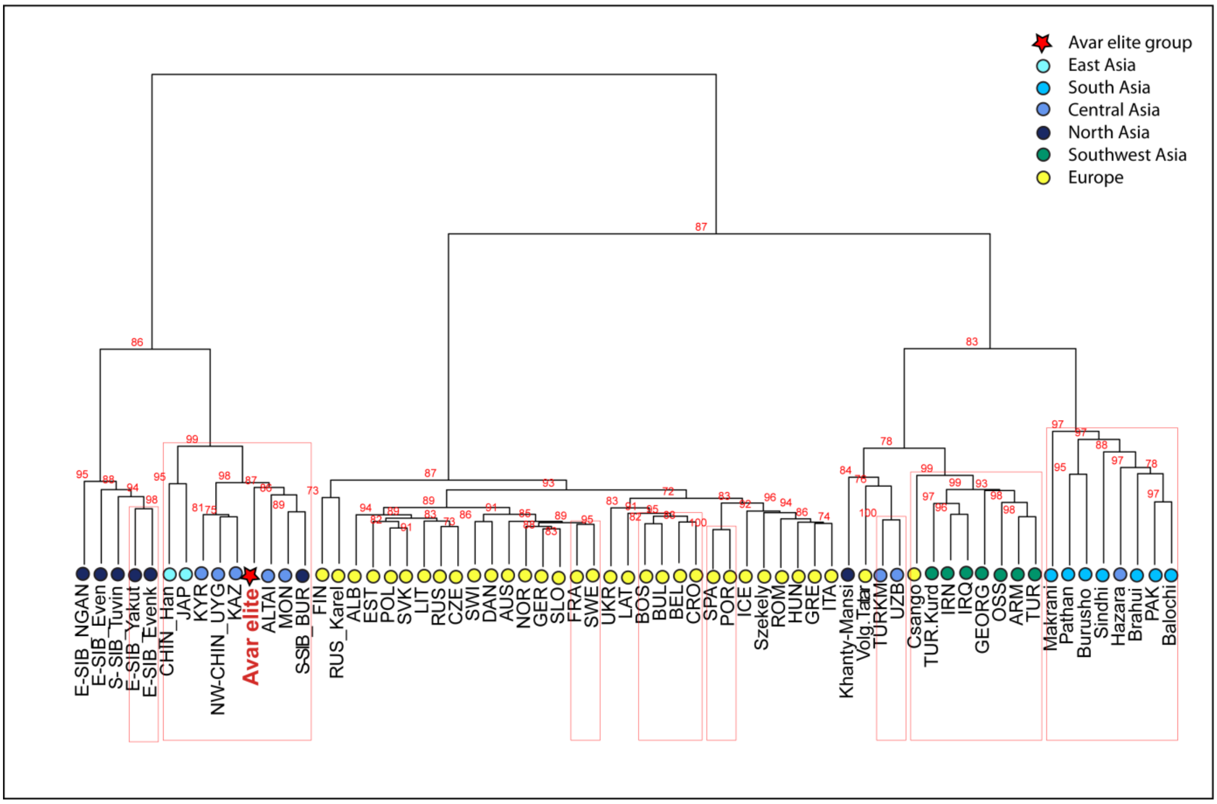

**Figure S8. Ward type clustering of the Avar elite group and 64 modern populations.** The Ward-type clustering shows the grouping of Asian, European, Central-South Asian and Near-Eastern and Caucasian populations under major branches of tree, and the Avars are situated on the Asian branch and together with Altaian (ALTAI), Mongolian (MON) and Transbaikalian Buryat (S-SIB_BUR) populations, furthermore with populations of Kyrgyzstan (KYR), Uyghurs from northwest China (NW-CHIN_UYG) and Kazakhstan (KAZ), and Chinese (CHIN) and Japanese (JAP). P values in percent are given as red numbers on the dendogram, where red rectangles indicate clusters with significant p values. The abbreviations and references are presented in Table S3.

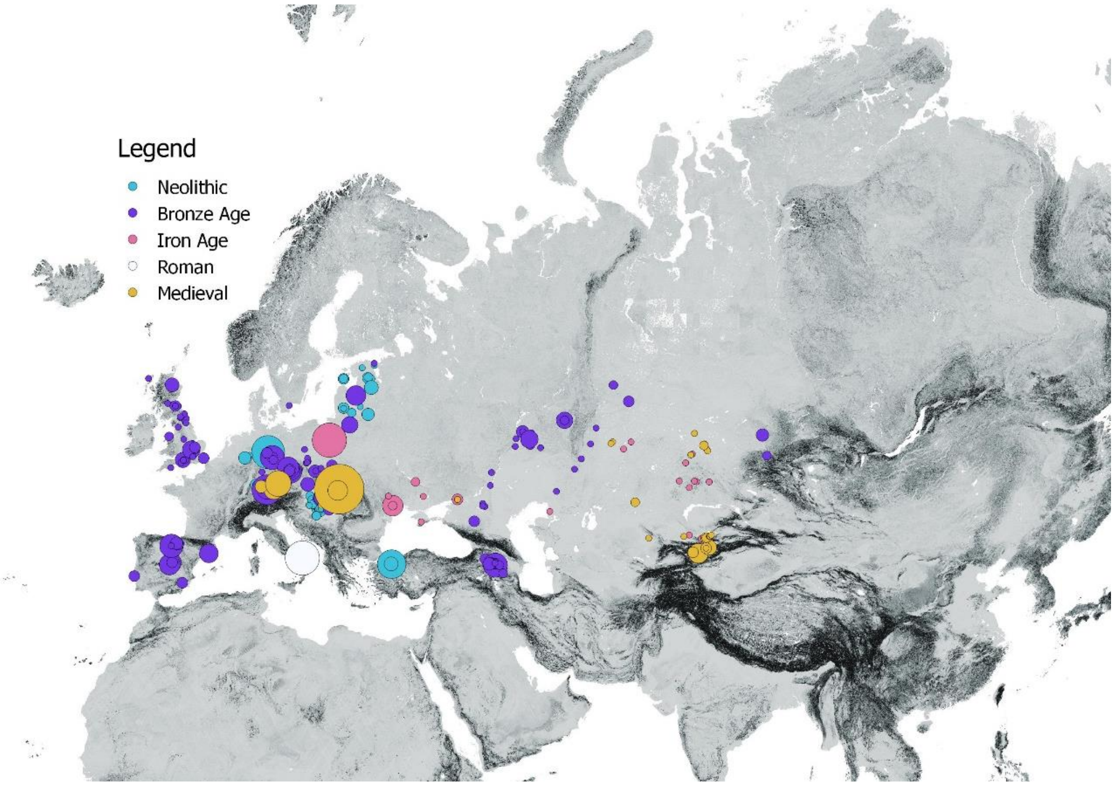

**Figure S9. Map of the ancient reference samples (only full mitogenomes) used in the $F_{ST}$ and MDS analyses.** References are seen in Table S4 and S8. Map was drawn by open source QGIS (Quantum GIS) software v. 2.8.1.

**Figure S10a-o: Phylogenetic trees** (For all of the phylogenetic trees the abbreviations and references are summarised in Table S6)

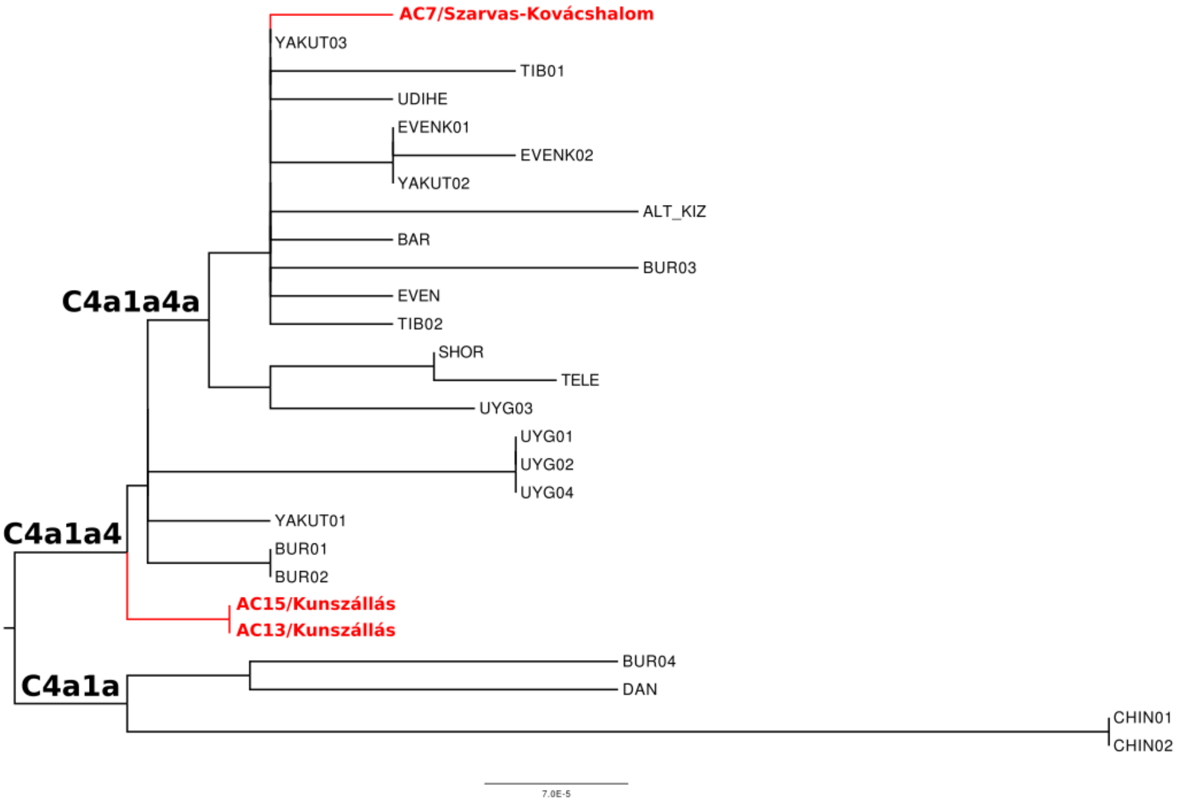

**Figure S10a. Phylogenetic tree of C4a1a haplogroup.**

This phylogenetic tree contains mostly North and East-Central Asian individuals. Samples AC13 and AC15 share identical haplotypes which may refer to close kinship.

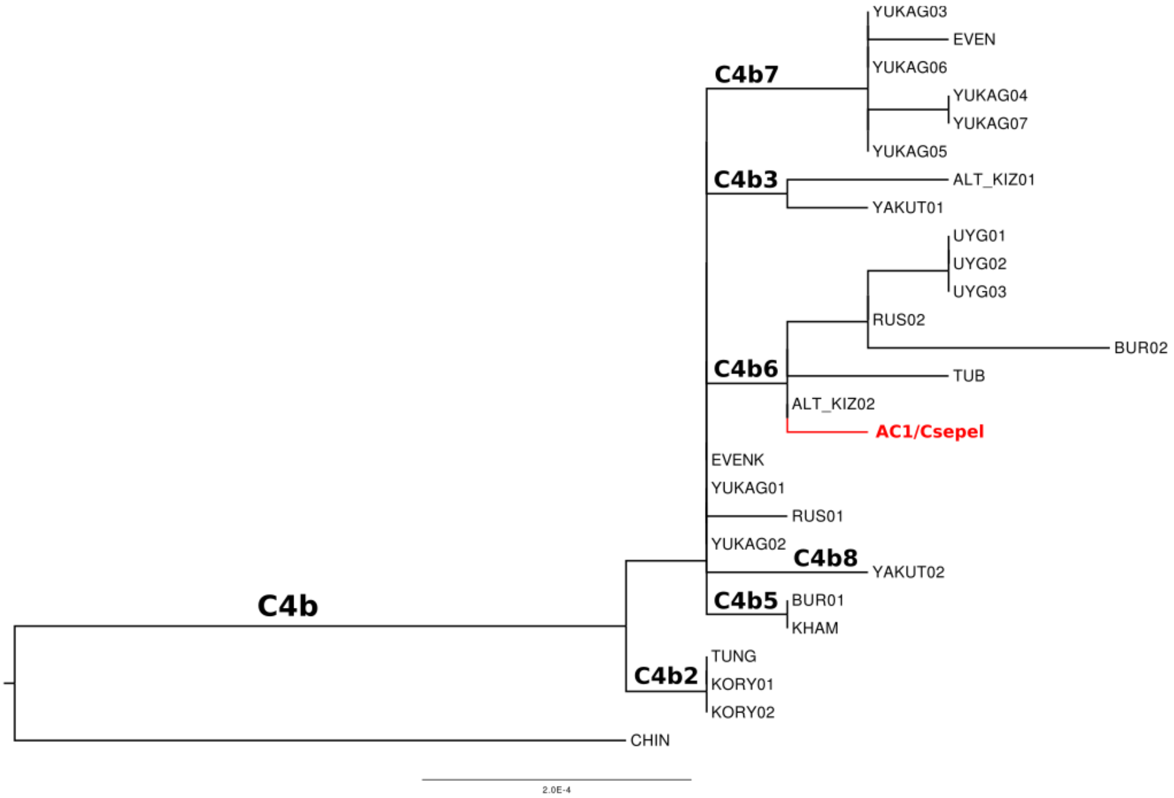

**Figure S10b. Phylogenetic tree of C4b haplogroup.**
This tree displays solely North – and East-Central Asians.

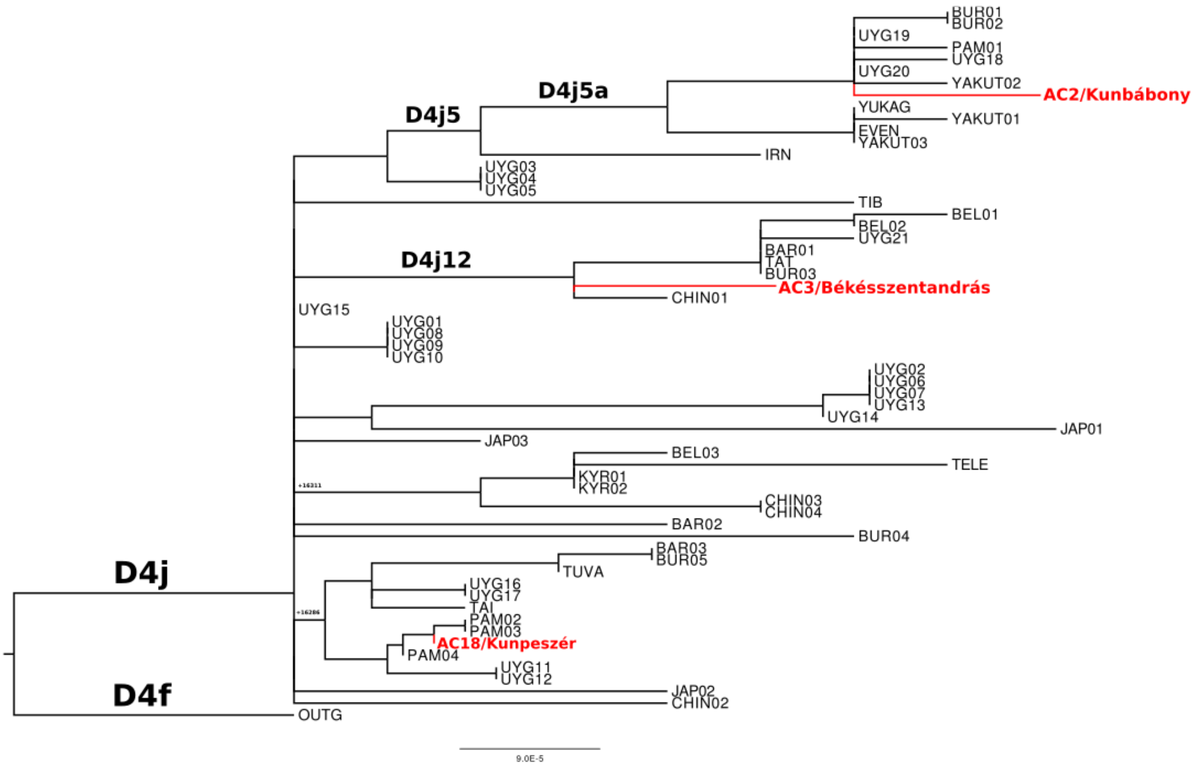

**Figure S10c. Phylogenetic tree of D4j haplogroup.**

Uyghur dominance on this tree is conspicuous, however the phylogeographic origin of the samples are widely distributed.

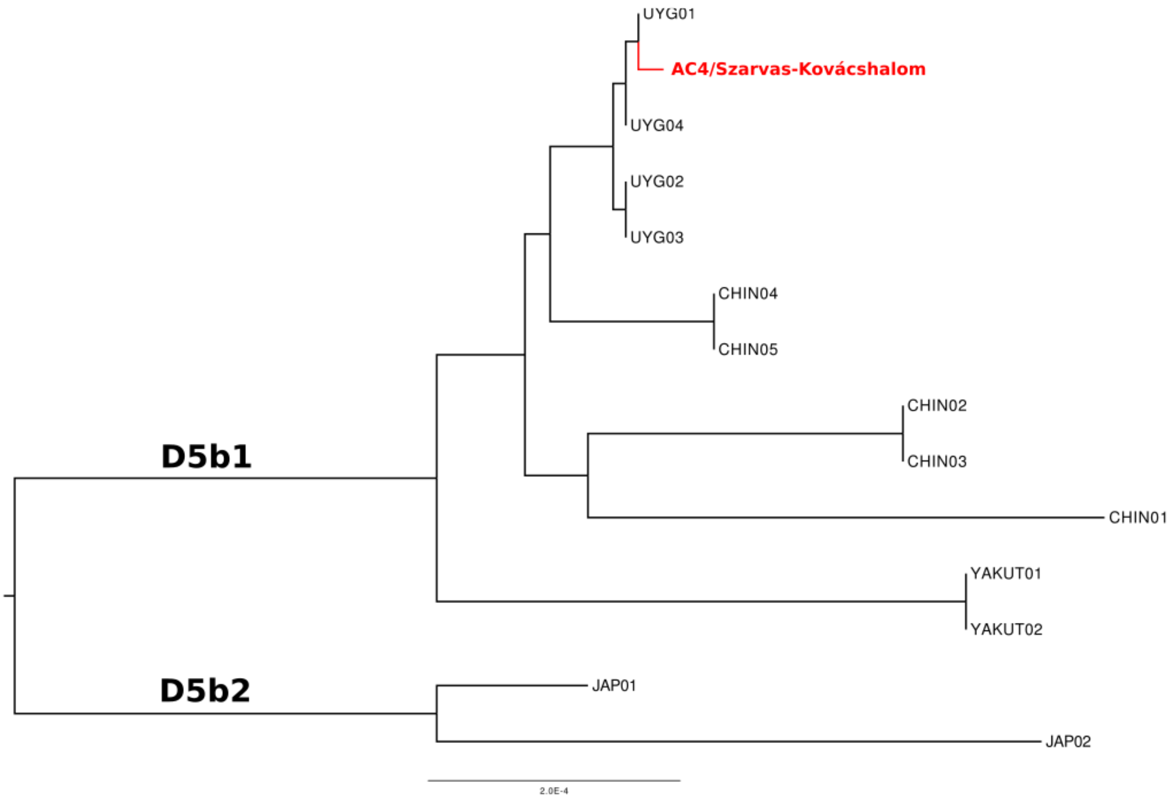

**Figure S10d. Phylogenetic tree of D5b haplogroup.**
Phylogeographically clearly distributed phylogenetic tree of this subhaplogroup.

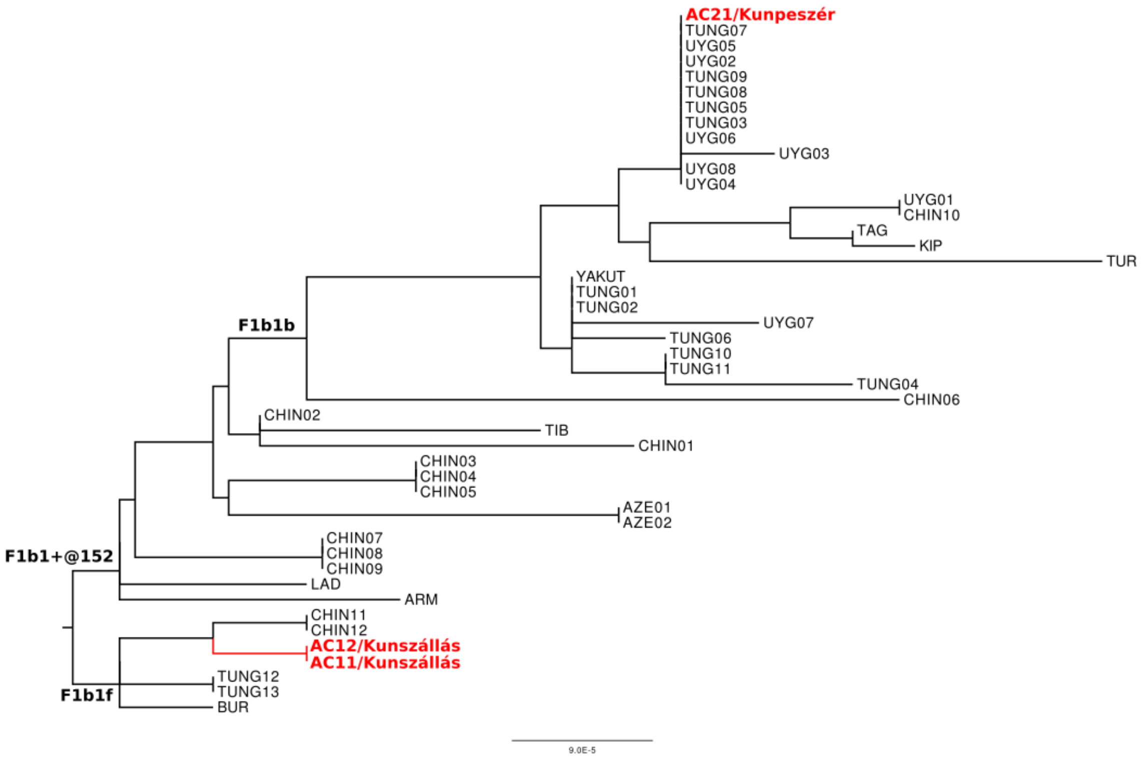

**Figure S10e. Phylogenetic tree of F1b1 haplogroup.**
Phylogeographically roughly resolved tree, where AC11 and AC12 share identical haplotypes with each other as well AC21 with Tungusic and Uyghur individuals.

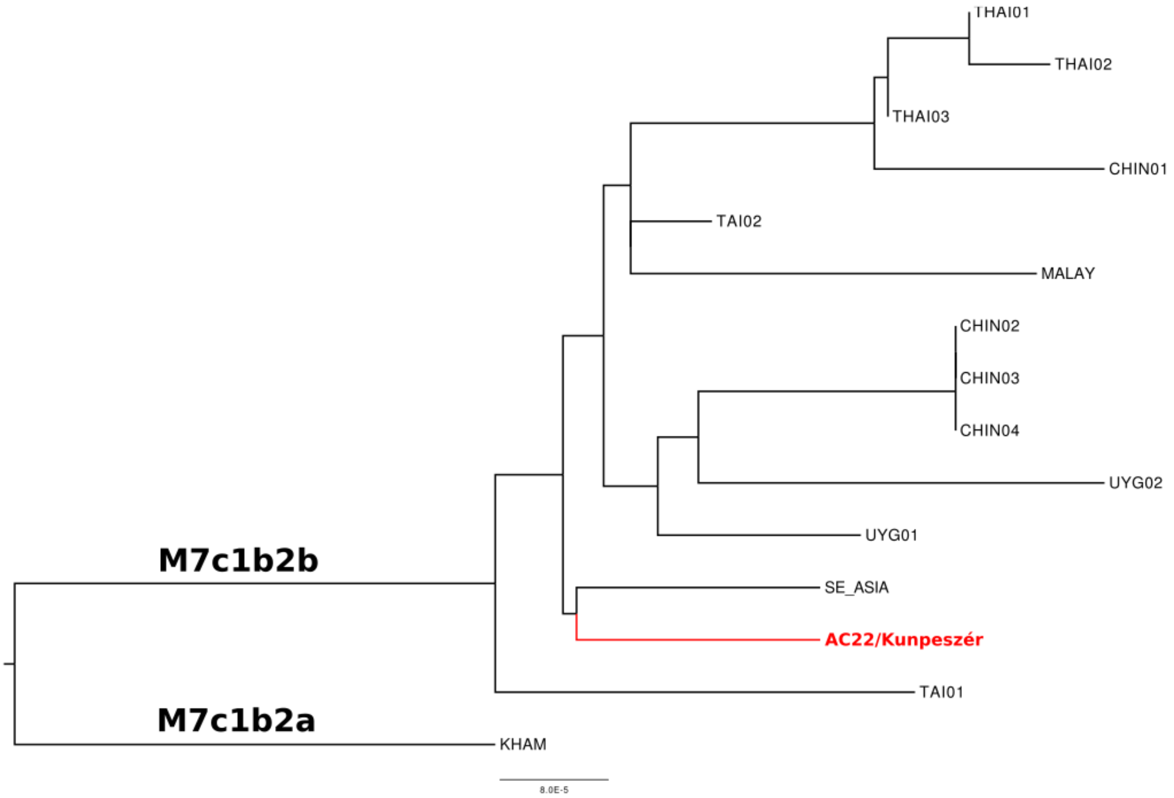

**Figure S10f. Phylogenetic tree of M7c1b2 haplogroup.**
Phylogeographically roughly resolved tree, where AC22 shows a rather South-East Asian origin, but due to the low sample number, closer relationship with Uyghurs cannot be rejected.

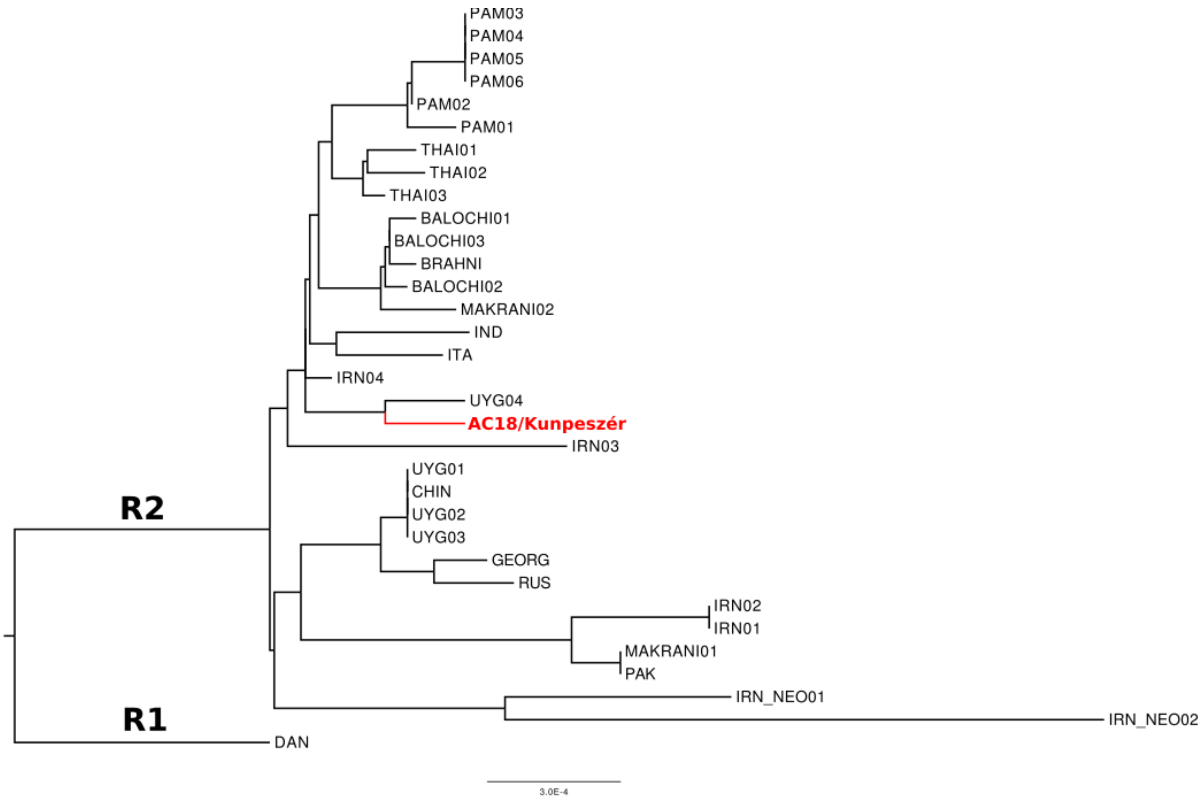

**Figure S10g. Phylogenetic tree of R2 haplogroup.**

This tree shows a rather South-Central Asian distribution of individuals, where AC18 still cluster together with a Uyghur.

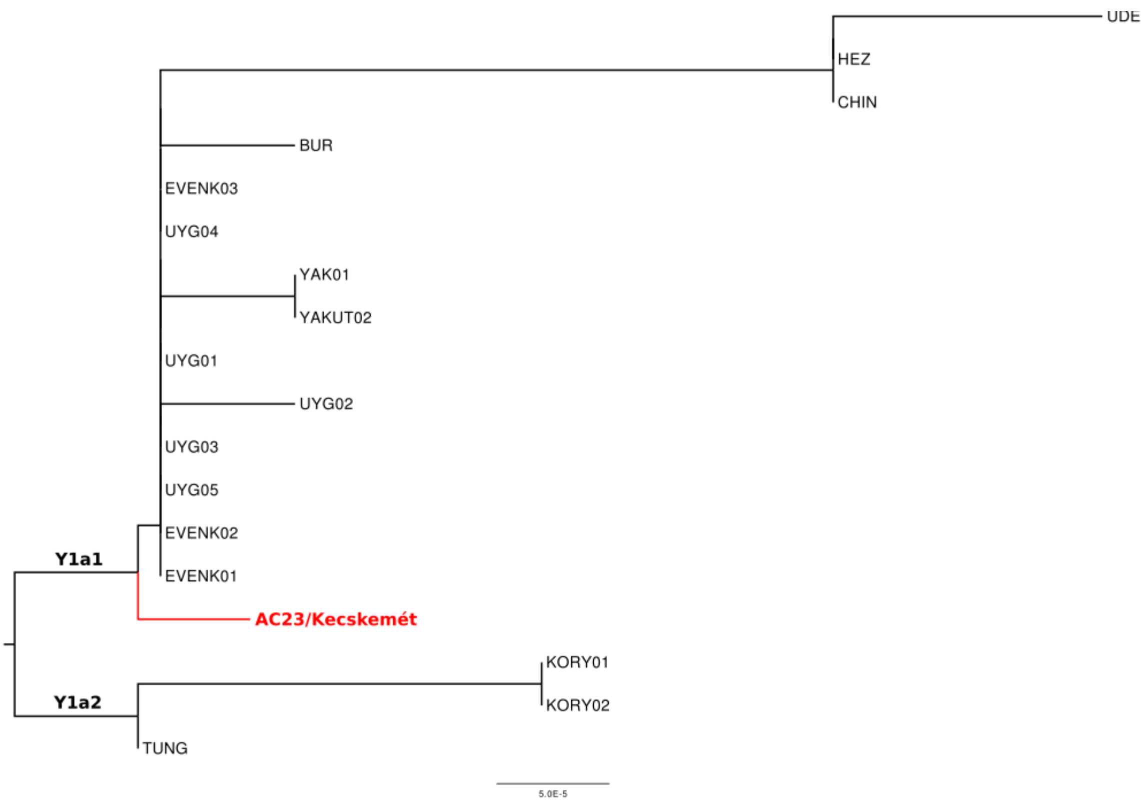

**Figure S10h. Phylogenetic tree of Y1a haplogroup.**

Rather North-East Asian distribution of individuals, the many identical haplotypes and short branch lengths may indicate a relatively young age for this subhaplogroup.

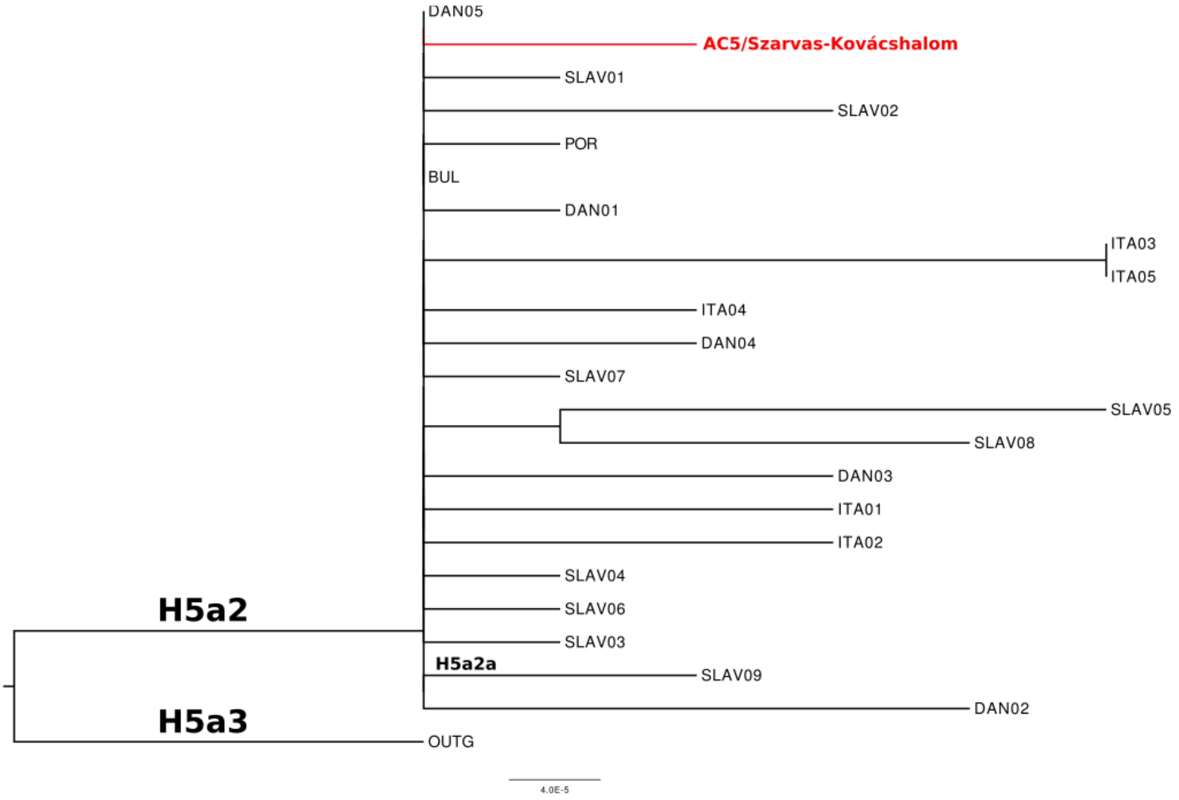

**Figure S10i. Phylogenetic tree of H5a haplogroup.**
This tree consists of European individuals with a dominance of northern Slavs.

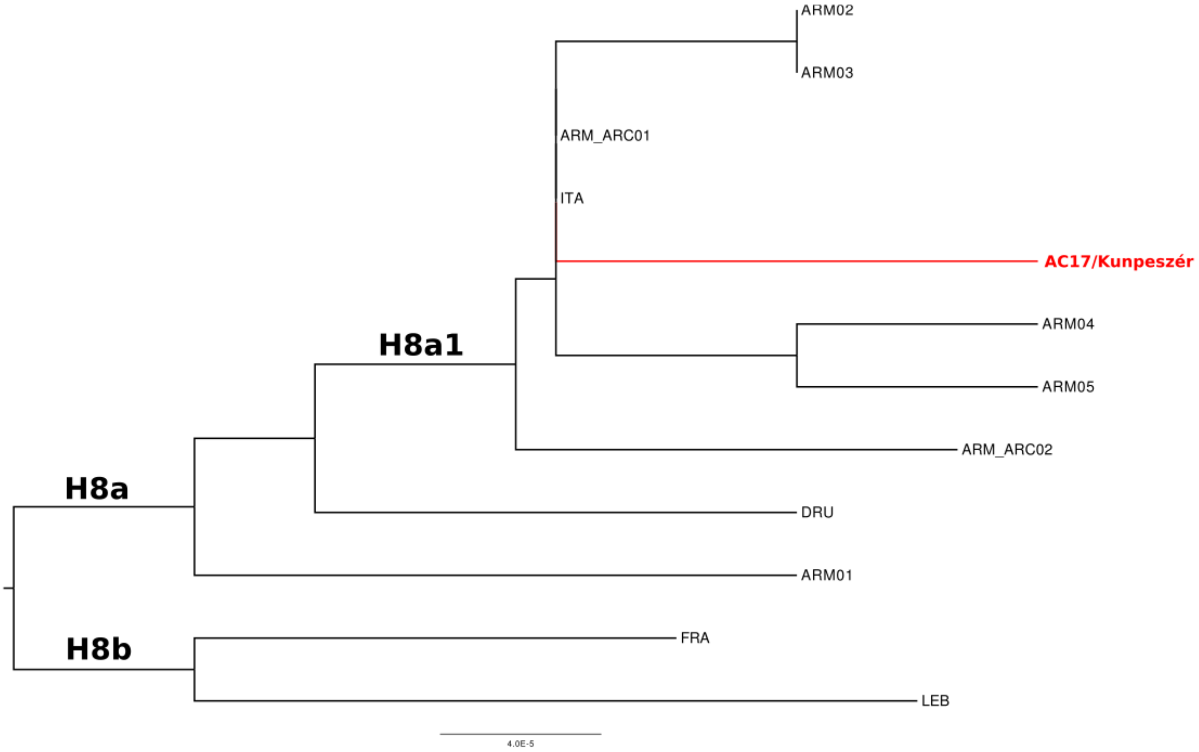

**Figure S10j. Phylogenetic tree of H8 haplogroup.**

The tree consists of mainly Caucasian (Armenian) individuals, where AC17 is well nested monophyletically to them.

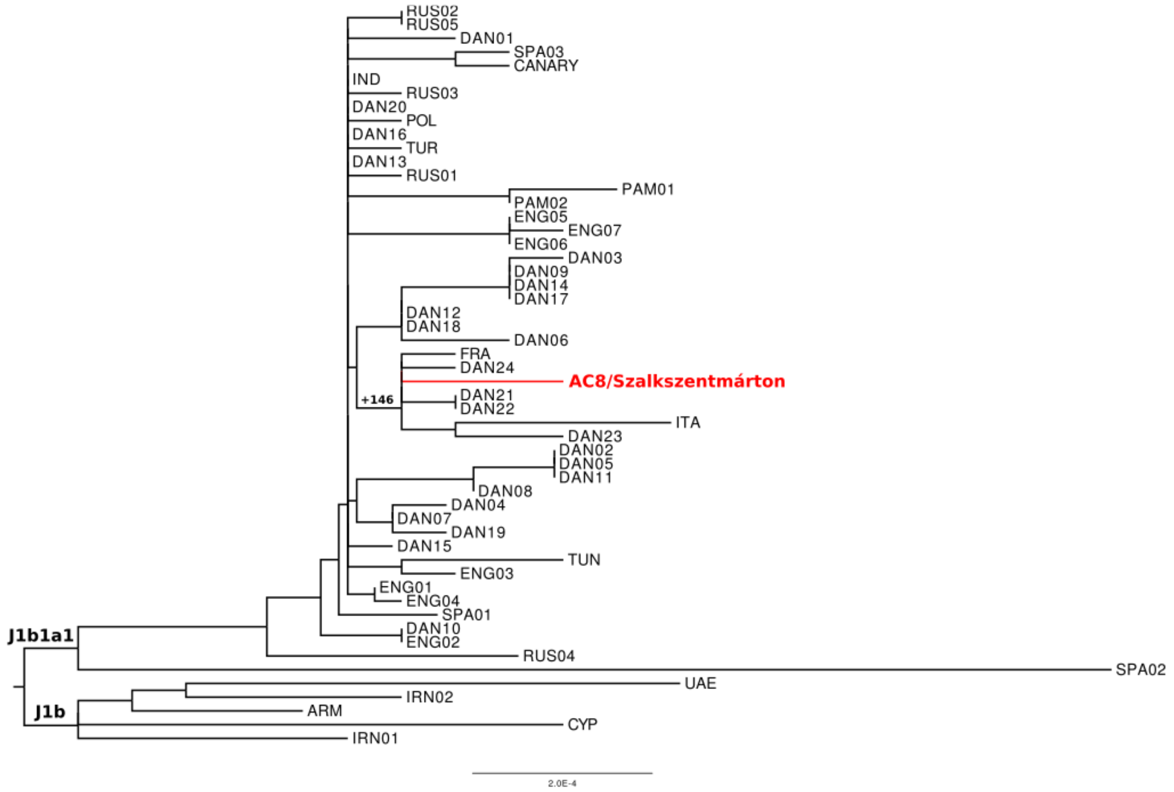

**Figure S10k. Phylogenetic tree of J1b haplogroup.**
This phylogenetic tree is of a typical European character, where more precise phylogeographic distribution cannot be inferred.

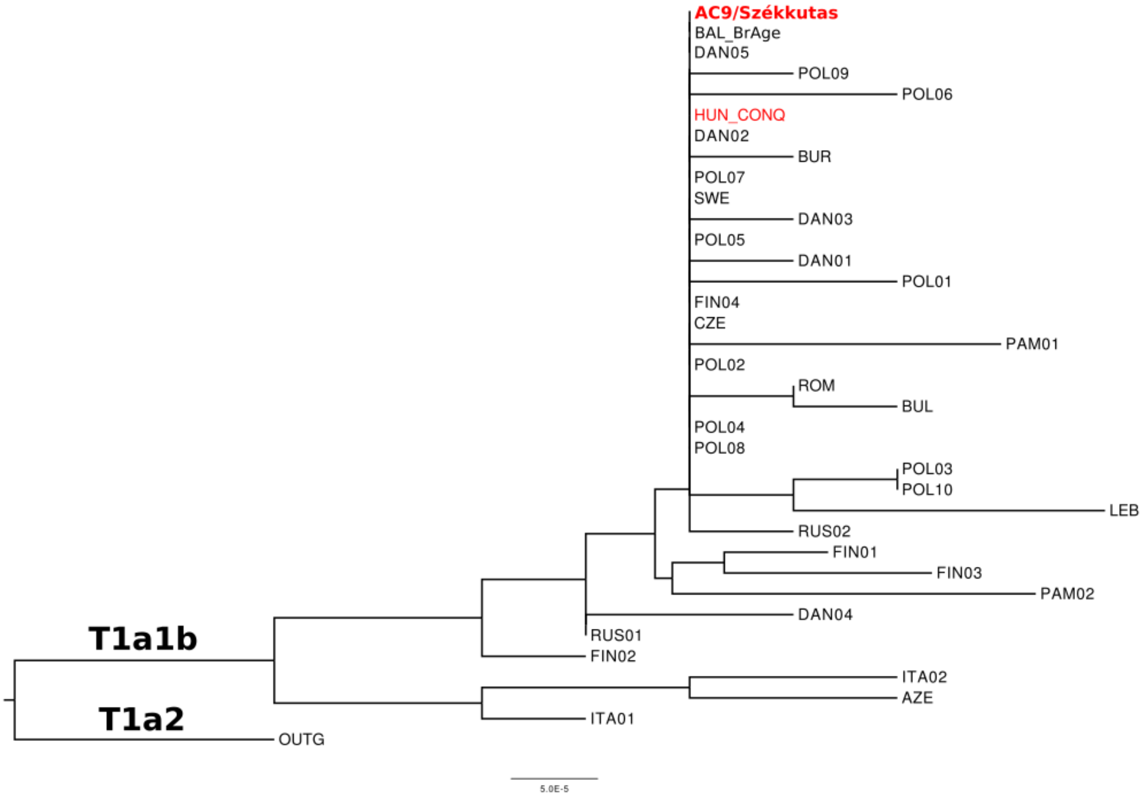

**Figure S10l. Phylogenetic tree of T1a1b haplogroup.**

The T1a1b tree represents mostly European populations. Interestingly, the investigated Avar sample AC9 has identical sequence to a Hungarian conqueror from Karos cemetery. However, this is the first and only observance of such relation, and according to the other 11 identical individuals from various regions, this may indicate that, at least for the Avar elite's maternal line, this connection is rather incidental than direct.

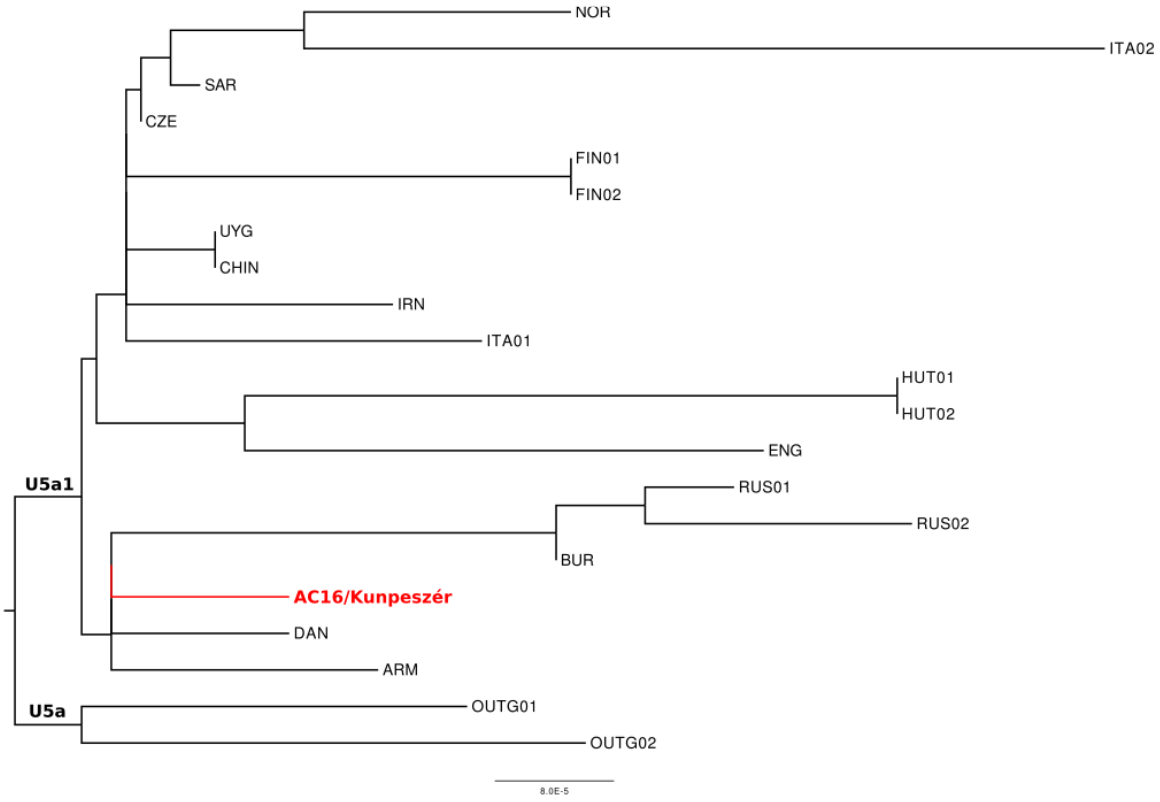

**Figure S10m. Phylogenetic tree of U5a haplogroup.**
This haplogroup is phylogeographically unresolvable due to its deep branching.

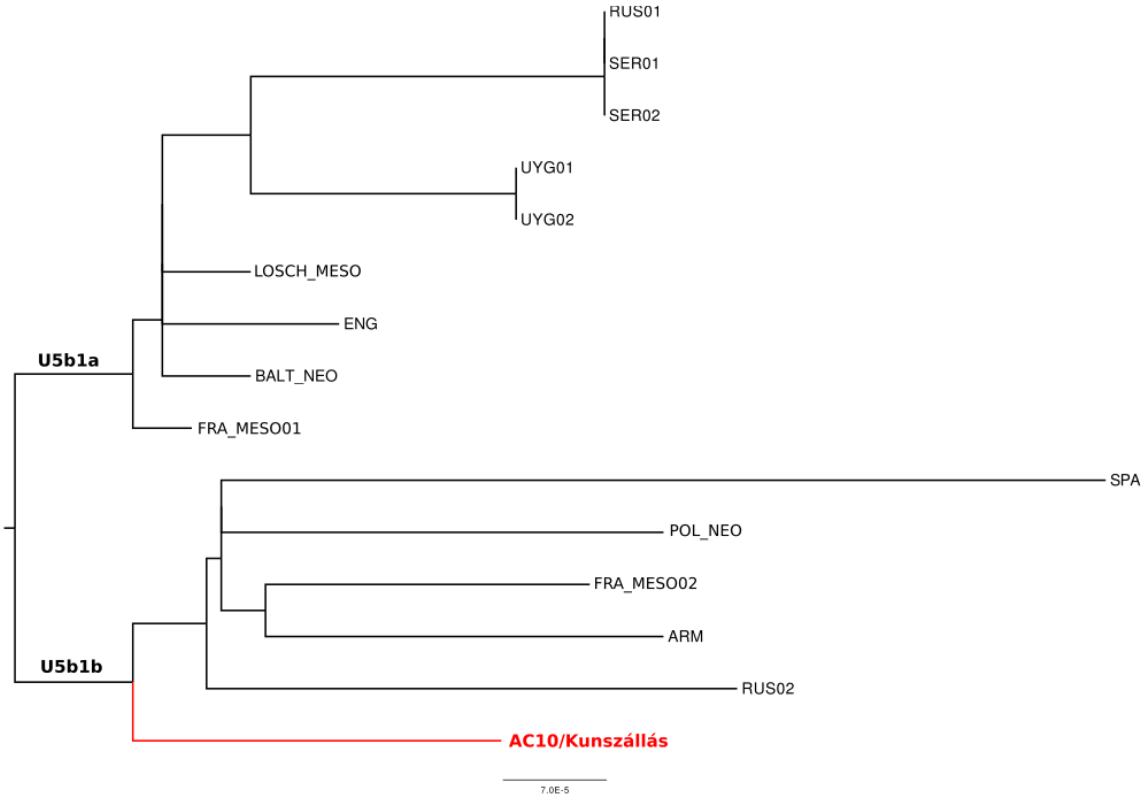

**Figure S10n. Phylogenetic tree of U5b1 haplogroup.**

The basal position of AC10 in subhaplogroup and wide and deep origins of samples make this tree phylogeographically unresolvable.

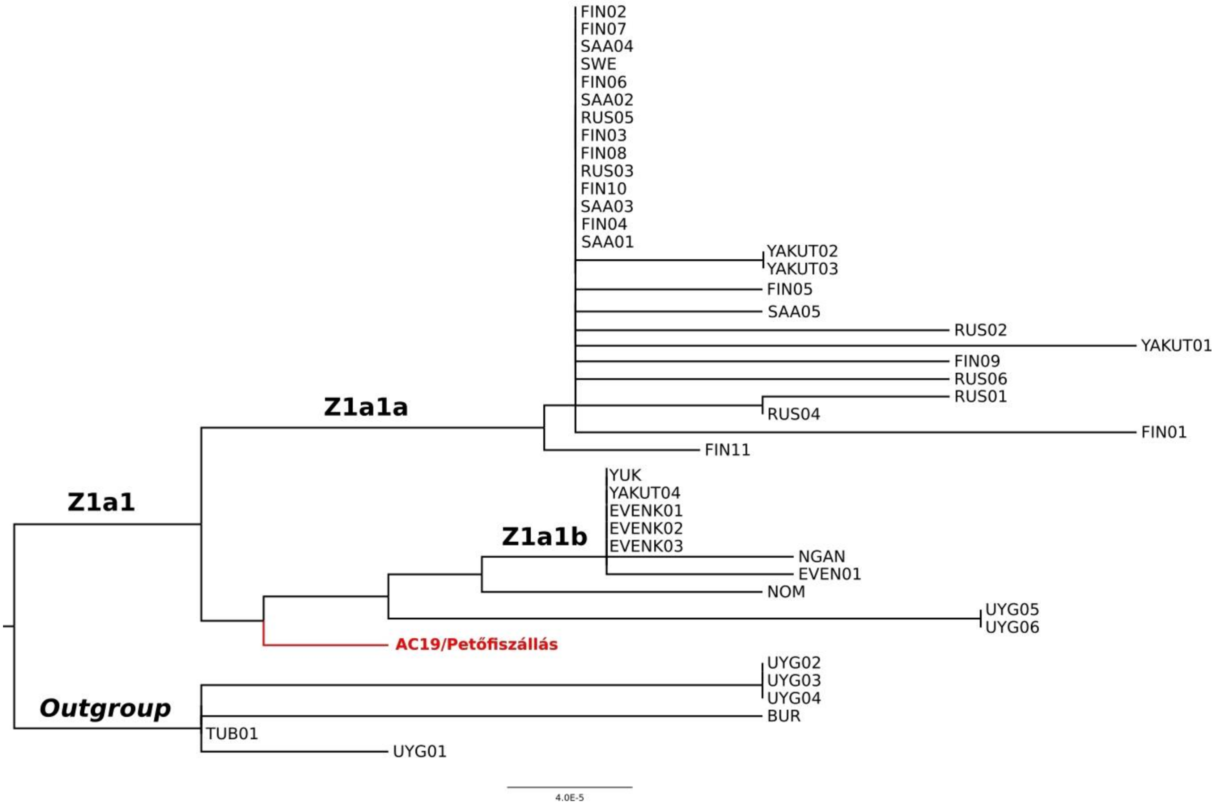

**Figure S10o. Phylogenetic tree of Z1a1 haplogroup.**

The individuals of Z1a1 subhaplogroup show their basal position to Z1a1b, where AC19 is the most basal to all of them. Interestingly, while Z1a1a is also the direct descendant of the Z1a1 subhaplogroup, it has no direct ancestry among them and possesses more Western Eurasian characteristics, than the rest of the tree. Six individuals from the Z1a subgroup were used as outgroup in the visualization.

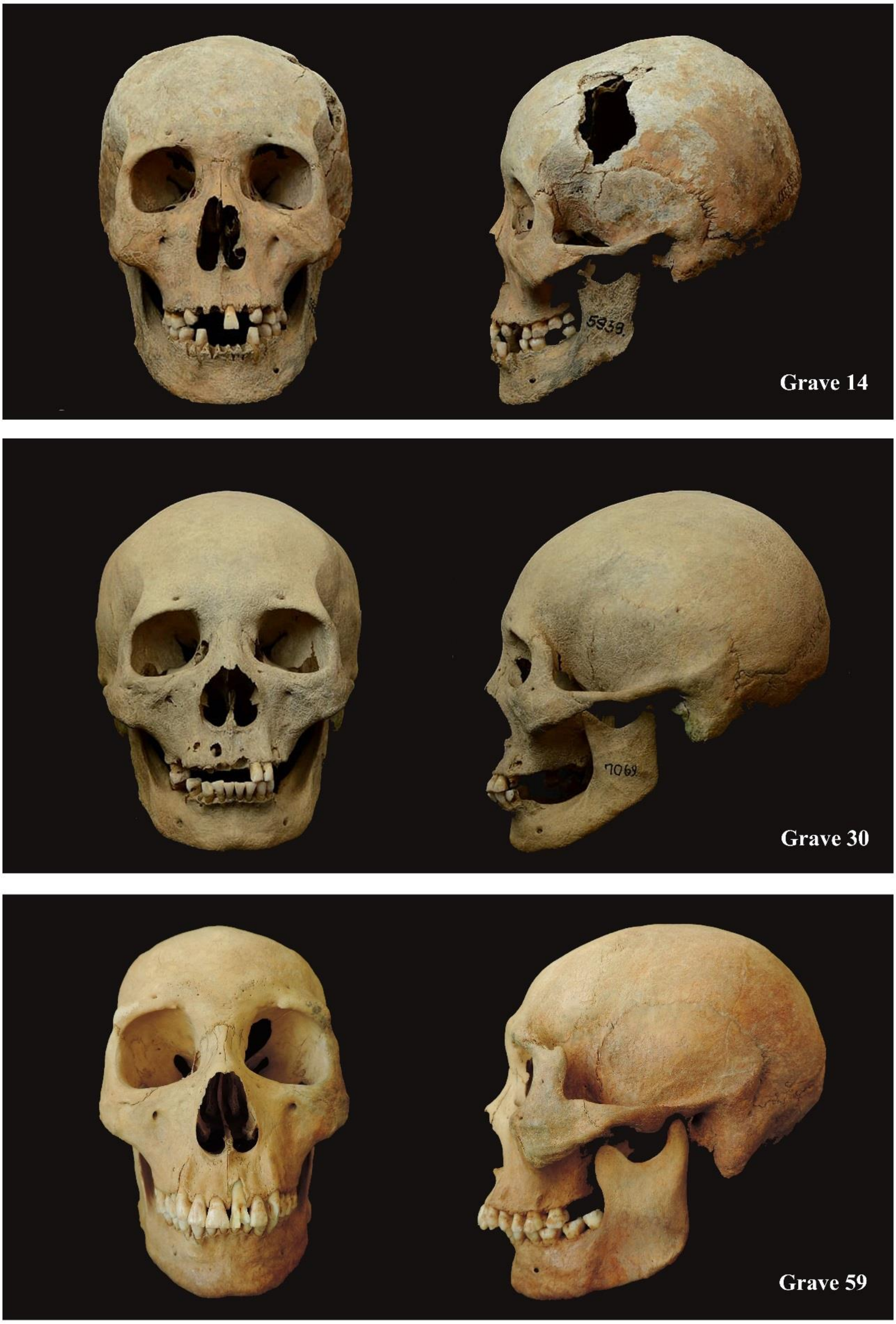

**Figure S11. Characteristic Avar skulls from the cemetery of Kunszállás (Hungary).** Photos were taken by Luca Kiss and Erika Molnár.
